# Single-Cell Transcriptomic Analysis of Human Lung Reveals Complex Multicellular Changes During Pulmonary Fibrosis

**DOI:** 10.1101/296608

**Authors:** Paul A. Reyfman, James M. Walter, Nikita Joshi, Kishore R. Anekalla, Alexandra C. McQuattie-Pimentel, Stephen Chiu, Ramiro Fernandez, Mahzad Akbarpour, Ching-I Chen, Ziyou Ren, Rohan Verma, Hiam Abdala-Valencia, Kiwon Nam, Monica Chi, SeungHye Han, Francisco J. Gonzalez-Gonzalez, Saul Soberanes, Satoshi Watanabe, Kinola J.N. Williams, Annette S Flozak, Trevor T. Nicholson, Vince K. Morgan, Cara L. Hrusch, Robert D. Guzy, Catherine A. Bonham, Anne I. Sperling, Remzi Bag, Robert B. Hamanaka, Gökhan M. Mutlu, Anjana V. Yeldandi, Stacy A. Marshall, Ali Shilatifard, Luis A.N. Amaral, Harris Perlman, Jacob I. Sznajder, Deborah R. Winter, Monique Hinchcliff, A. Christine Argento, Colin T. Gillespie, Jane D’Amico Dematte, Manu Jain, Benjamin D. Singer, Karen M. Ridge, Cara J. Gottardi, Anna P. Lam, Ankit Bharat, Sangeeta M. Bhorade, G.R. Scott Budinger, Alexander V. Misharin

**Affiliations:** Northwestern University, Feinberg School of Medicine, Department of Medicine, Division of Pulmonary and Critical Care Medicine, Chicago, IL 60611, USA; Northwestern University, Feinberg School of Medicine, Department of Surgery, Division of Thoracic Surgery, Chicago, IL, 60611, USA; Northwestern University, Feinberg School of Medicine, Department of Medicine, Division of Rheumatology, Chicago, IL 60611, USA; University of Chicago, Section of Pulmonary and Critical Care Medicine, Chicago, IL, 60637; Northwestern University, Feinberg School of Medicine, Department of Pathology, Chicago, IL 60611, USA; Northwestern University, Feinberg School of Medicine, Department of Biochemistry and Molecular Genetics, Chicago, IL 60611, USA; Northwestern University, Weinberg College of Arts and Sciences, Department of Chemical and Biological Engineering Evanston, IL 60201, USA

## Abstract

Pulmonary fibrosis is a devastating disorder that results in the progressive replacement of normal lung tissue with fibrotic scar. Available therapies slow disease progression, but most patients go on to die or require lung transplantation. Single-cell RNA-seq is a powerful tool that can reveal cellular identity via analysis of the transcriptome, but its ability to provide biologically or clinically meaningful insights in a disease context is largely unexplored. Accordingly, we performed single-cell RNA-seq on lung tissue obtained from eight transplant donors and eight recipients with pulmonary fibrosis and one bronchoscopic cryobiospy sample. Integrated single-cell transcriptomic analysis of donors and patients with pulmonary fibrosis identified the emergence of distinct populations of epithelial cells and macrophages that were common to all patients with lung fibrosis. Analysis of transcripts in the Wnt pathway suggested that within the same cell type, Wnt secretion and response are restricted to distinct non-overlapping cells, which was confirmed using *in situ* RNA hybridization. Single-cell RNA-seq revealed heterogeneity within alveolar macrophages from individual patients, which was confirmed by immunohistochemistry. These results support the feasibility of discovery-based approaches applying next generation sequencing technologies to clinically obtained samples with a goal of developing personalized therapies.

**One Sentence Summary:** Single-cell RNA-seq applied to tissue from diseased and donor lungs and a living patient with pulmonary fibrosis identifies cell type-specific disease-associated molecular pathways.

## Introduction

Pulmonary fibrosis is defined as the progressive replacement of alveolar tissue with fibrotic scar that threatens alveolar gas exchange and reduces lung compliance. The resulting increased work of breathing and hypoxemia cause progressive respiratory failure and eventual death. While the cause of pulmonary fibrosis is often not identified (idiopathic pulmonary fibrosis, IPF), pulmonary fibrosis can be attributed to connective tissue disease, environmental exposures (hypersensitivity pneumonitis), occupational exposures (silicosis, asbestosis) or drugs in some patients. Diagnostic approaches to distinguish between these causes of pulmonary fibrosis, including surgical lung biopsy, remain imprecise, and there are few laboratory features that predict responsiveness to therapy (*1, 2*). The clinical and economic burden of pulmonary fibrosis is sizeable: the worldwide incidence of IPF is increasing and attributable medical costs for patients with IPF in the United States alone have been estimated at nearly 2 billion dollars (*3, 4*). Despite decades of research, only a handful of therapies are available. While these therapies slow disease progression, the response is variable, and lung transplantation represents the only option for temporary cure. Accordingly, there has been a call to leverage cutting-edge genomic techniques and systems biology approaches to provide molecular insights into pulmonary fibrosis that can be used to develop personalized approaches to diagnosis and therapy (*5*).

While every cell in the body has an identical genome, the quantity of mRNA molecules encoded by individual genes, collectively referred to as the transcriptome, differs widely as a function of cell type. Indeed, investigators have used next generation sequencing technology (RNA-seq) to develop an organ-level transcriptomic map of the human body (*6*). The lung is a complex tissue comprised of more than forty cell populations (*7*). Because each of these populations has a distinct transcriptome, previous investigations of the “lung transcriptome” during fibrosis using whole lung tissue were limited by changes in the cellular composition of the lung during disease, which could mask relevant changes in individual cell populations (*8–11*). Transcriptomic analysis of flow cytometry-sorted cell populations from the lung provides an alternative, but requires *a priori* assumptions about cell surface markers whose expression may change during disease. The advent of single-cell RNA-seq allows reliable identification of even closely-related cell populations (e.g. differentiating between myeloid cell populations in the bone marrow) (*12*).

Single-cell RNA-seq methods also allow for the measurement of abundantly expressed transcripts in cell populations for which there are no reliable surface markers, and provide the opportunity to assess heterogeneity of gene expression in individual lung cell populations during health and disease (*13, 14*).

Here, we used droplet-based single-cell RNA-seq to analyze lung tissue from patients with pulmonary fibrosis and lung transplant donors. We identified multiple cell populations in the normal lung, and transcriptionally distinct macrophage and epithelial cell populations in patients with pulmonary fibrosis. Differentially expressed transcripts between patients with pulmonary fibrosis and donors in each cell population were enriched for genes previously associated with pulmonary fibrosis. Analysis of transcripts in the Wnt pathway suggested restriction of Wnt secretion and Wnt response to distinct non-overlapping cell populations within the same cell type. This was confirmed using *in situ* RNA hybridization. Single-cell RNA-seq accurately characterized the expression of different fibrosis-related proteins in alveolar macrophages from individual patients, and was useful for the identification of rare cell populations including senescent cells. These results support the feasibility of discovery-based approaches applying next generation sequencing technologies to clinically obtained samples with a goal of developing personalized therapies.

## Results

### Single-cell RNA-seq identifies multiple cell populations in the mouse lung

To determine whether analysis of single-cell RNA-seq data could resolve cellular diversity in the normal lung, we analyzed single-cell suspensions generated from two mouse lungs. To better identify rare cell populations, we combined data from both mice, analyzing 14,514 total cells.

Clustering was performed using K-nearest neighbor-based clustering using Euclidean distance in principal component analysis space as implemented in Seurat R toolkit (*15, 16*). Analysis identified 23 individual clusters (Figure 1A) and cell identity was assigned based on expression of marker genes (Figure 1B). We identified all well-known cell types including alveolar type II cells, ciliated and club airway epithelial cells, alveolar macrophages and dendritic cells. We were also able to identify rare and difficult to isolate cell populations, including alveolar type I cells, several types of endothelial and lymphatic cells, megakaryocytes, innate lymphoid cells and mesothelial cells (Figure 1B, Supplemental Table 1). All cell types were well represented in both libraries, indicating sampling reproducibility (Supplementary Figure S1). This dataset can be explored using the Seurat package in R or the interactive Loupe Cell Browser (Supplemental Files 1 and 2)(*15, 16*).

**Figure 1.**
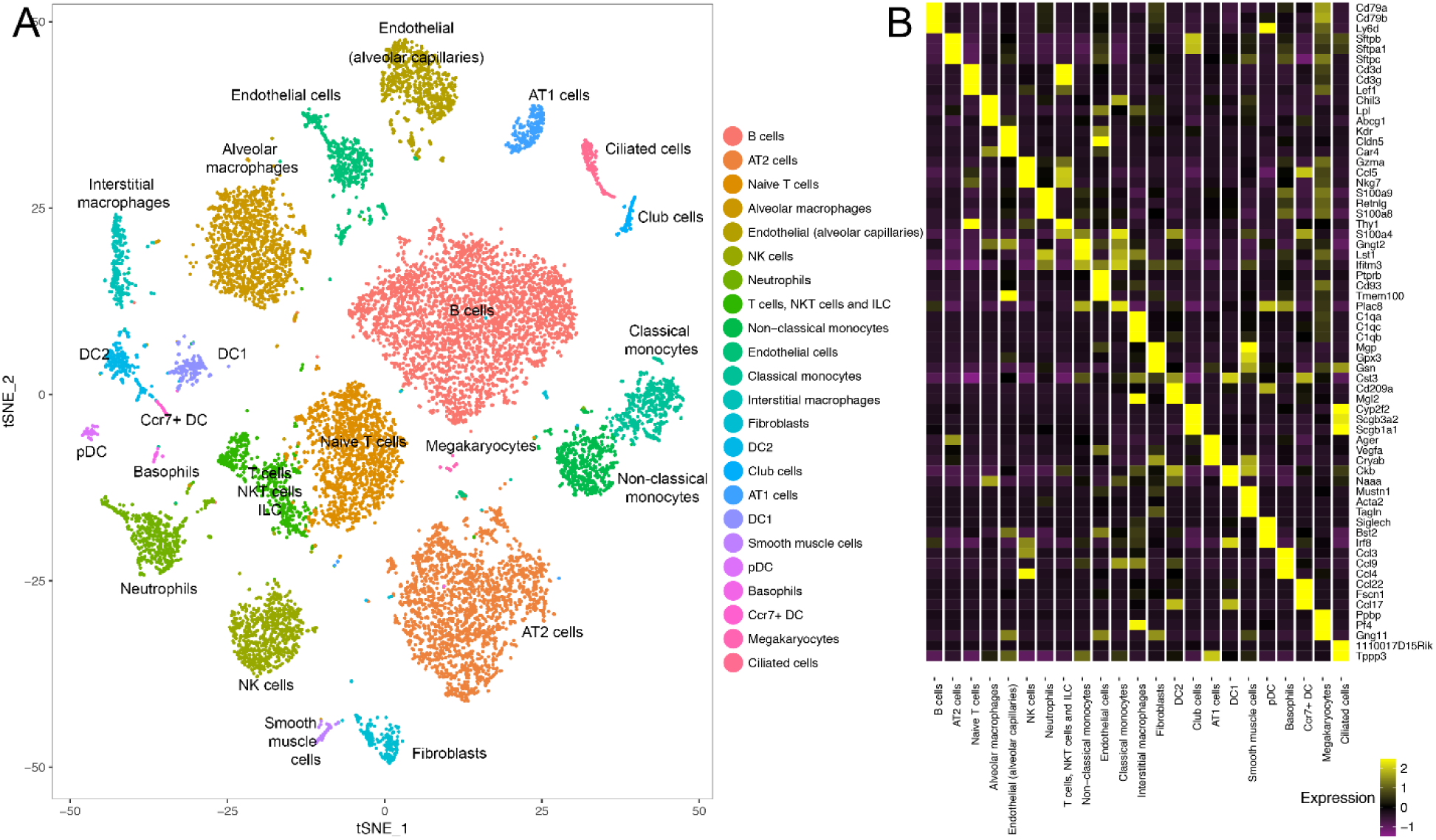
Single-cell RNA-seq identified 23 cell populations in the normal mouse lung, including several rare populations. A. t-distributed stochastic neighbor embedding plot shows clustering of 14,514 cells, data from 2 mice. B. Heatmap shows average expression per cell type of the top 3 marker genes for each cell type.

### Study Population

Single-cell RNA-seq was performed on eight donor lung biopsies and eight patients with pulmonary fibrosis attributed to IPF (4 patients), systemic sclerosis (2 patients), polymyositis (1 patient) and chronic hypersensitivity pneumonitis (1 patient). All samples were obtained at the time of transplantation. Separately, we performed single-cell RNA-seq using one bronchoscopic cryobiopsy sample from a patient subsequently diagnosed with IPF. Bulk-cell RNA-seq was performed on samples of lung tissue obtained from 14 donors prior to transplantation and 8 transplant recipients with pulmonary fibrosis. The median age of patients with pulmonary fibrosis was 56.0 years (interquartile range ± 14.5 years). Eight (47.0%) were male and 6 (35.3%) were former smokers. Characteristics of patients with pulmonary fibrosis are reported in Table 1 and characteristics of donors are reported in Table 2.

**Table 1.** Characteristics of Patients with Pulmonary Fibrosis. *Transplant recipients with pulmonary fibrosis included in single-cell RNA-seq analysis. †Patient with IPF undergoing cryobiopsy included in single-cell RNA-seq analysis. ‡Transplant recipients with pulmonary fibrosis included in bulk-cell RNA-seq analysis. (ILD, Interstitial lung disease; IPF, idiopathic pulmonary fibrosis)

| Age | Sex | Diagnosis | History of Smoking | Supplemental Oxygen Use Prior to Transplant |
| --- | --- | --- | --- | --- |
| 66* | Male | IPF | Yes | Yes |
| 60* | Male | IPF | No | Yes |
| 68* | Male | IPF | Yes | Yes |
| 72* | Female | IPF | No | Yes |
| 55* | Female | Hypersensitivity Pneumonitis | No | Yes |
| 39* | Female | Systemic sclerosis-associated ILD | No | Yes |
| 53* | Female | Systemic sclerosis-associated ILD | No | Yes |
| 37* | Female | Myositis-associated ILD | No | Yes |
| 71† | Female | IPF | Yes | No |
| 42‡ | Male | Systemic sclerosis-associated ILD | No | Yes |
| 65‡ | Male | IPF | No | Yes |
| 52‡ | Male | Systemic sclerosis-associated ILD | Yes | Yes |
| 52‡ | Female | Systemic sclerosis-associated ILD | No | Yes |
| 55‡ | Female | Myositis-associated ILD | No | Yes |
| 56‡ | Female | Hypersensitivity Pneumonitis | No | Yes |
| 60‡ | Male | Myositis-associated ILD | Yes | Yes |
| 67‡ | Male | IPF | Yes | No |

**Table 2.** Characteristics of lung transplant donors. *Used for single-cell RNA-seq analysis.

| Age (years) | Sex | Race | Cause of death |
| --- | --- | --- | --- |
| 41 | Male | Caucasian | Stroke |
| 57 | Male | Caucasian | Intracranial hemorrhage following fall |
| 64 | Male | African American | Intracranial hemorrhage |
| 49 | Female | Caucasian | Intracranial hemorrhage |
| 50 | Male | African American | Intracranial hemorrhage |
| 19 | Male | African American | Stroke following gunshot wound |
| 54 | Male | Caucasian | Intracranial hemorrhage following blunt trauma |
| 43 | Male | Hispanic | Anoxic brain injury following opiate overdose |
| 21 | Male | Caucasian | Intracranial hemorrhage following blunt trauma |
| 43 | Female | Caucasian | Anoxic brain injury |
| 50 | Male | Hispanic | Intracranial hemorrhage following blunt trauma |
| 51 | Male | African American | Stroke |
| 26 | Female | African American | Anoxic brain injury following opiate overdose |
| 40 | Male | Hispanic | Intracranial hemorrhage following blunt trauma |
| 28* | Male | Caucasian | Intracranial hemorrhage following blunt trauma |
| 55* | Male | Asian | Intracranial hemorrhage |
| 29* | Female | African American | Anoxic brain injury |
| 57* | Female | African American | Anoxic brain injury |
| 49* | Female | Caucasian | Intracranial hemorrhage |
| 22* | Female | African American | Anoxic brain injury following seizure |
| 21* | Male | African American | Head trauma from gunshot wound |
| 47* | Female | Caucasian | Intracranial Hemorrhage |

### Integrated analysis of single-cell RNA-seq data from healthy and diseased lungs identifies 13 cell populations

We performed single-cell RNA-seq on eight donor lungs, eight lungs explanted from patients with pulmonary fibrosis and one cryobiopsy from a patient with pulmonary fibrosis. An interactive file that includes these data is included in the online supplement (Supplementary File 3).

We reasoned that integrative analysis of multiple subjects might provide useful insights into the molecular events that drive pulmonary fibrosis. Since currently available computational tools for integrative analysis of multiple single-cell RNA-seq datasets did not perform well on very large and heterogeneous datasets, we limited our integrated analysis to the first 6 donor lungs and 6 lungs from patients with pulmonary fibrosis. In total, 48,937 cells were used for integrated analysis after filtering. We performed canonical correlation analysis (CCA) as implemented in the Seurat package and identified 13 clusters (*15*). A t-distributed stochastic neighbor embedding (t-SNE) plot with labeled cell types is shown in Figure 2A. Each cluster included cells from donors and patients with pulmonary fibrosis (Figure 2B). We assigned cell types to each cluster based on the expression of established markers (Figure 2C, D, Supplementary Figure S2, Supplementary Table 2).

**Figure 2.**
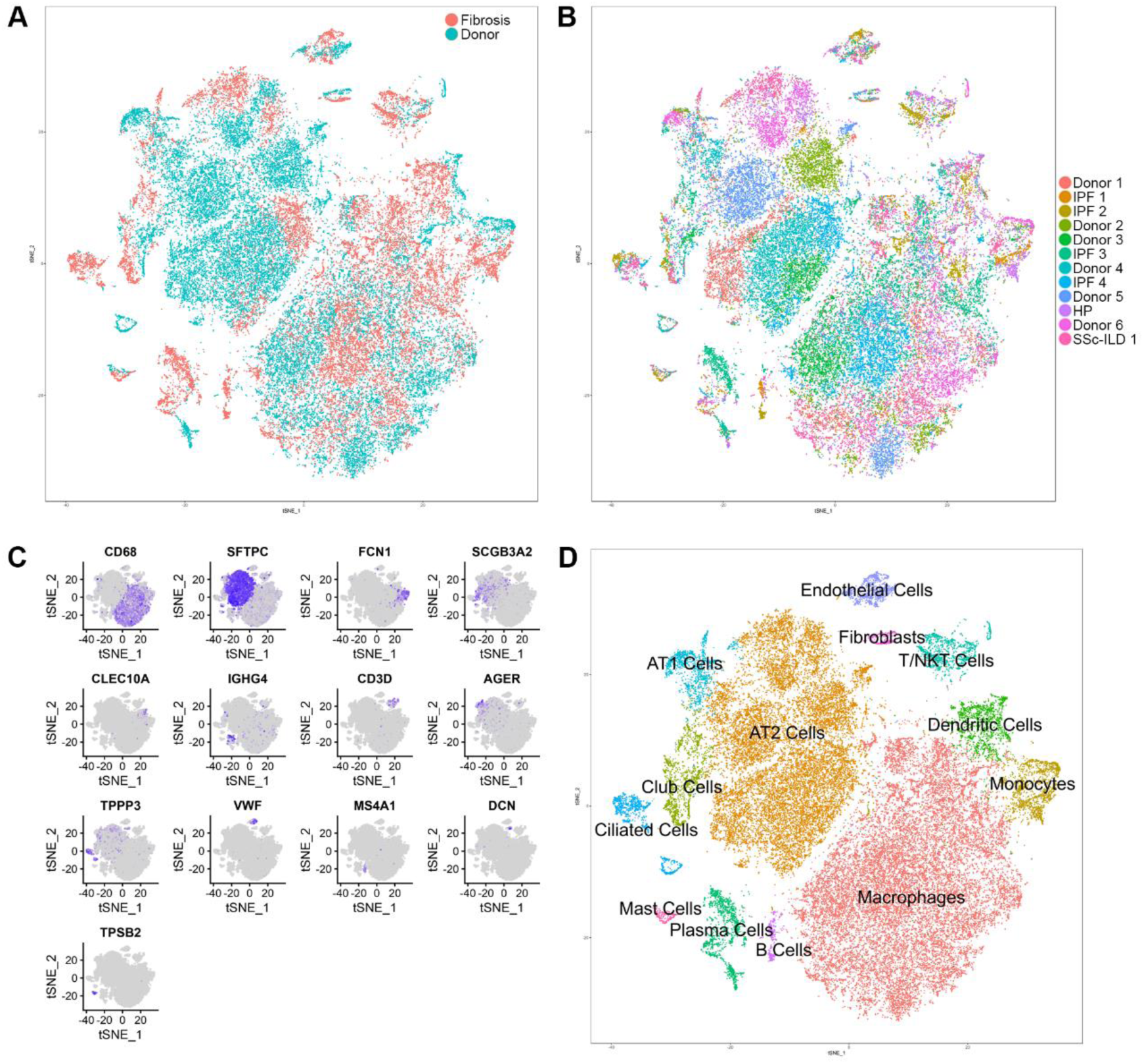
Integrated single-cell transcriptomic analysis of patients with pulmonary fibrosis identifies diverse lung cell populations. Droplet-based single-cell RNA-seq was performed on single-cell suspensions generated from six lung transplant donors and six lung transplant recipients with pulmonary fibrosis. All twelve samples were analyzed using the Seurat package with canonical correlation analysis procedure. Cells were clustered using a graph-based shared nearest neighbor clustering approach and visualized using a t-Distributed Stochastic Neighbor Embedding (t-SNE) plot. A. Cellular populations identified. B. Cells on the t-SNE plot of all 12 samples were colored according to patient group. Each population included cells from donors and patients with pulmonary fibrosis. C, D Canonical cell markers were used to label clusters by cell identity as represented in the t-SNE plot.

### Single-cell transcriptomic analysis reveals disease-associated changes in all identified lung cell populations

Comparison of gene expression within cell populations between samples from donor lungs and those from patients with pulmonary fibrosis identified significant differentially expressed genes in all 13 cell populations, including alveolar macrophages, alveolar type II cells, and fibroblasts (Figures 3A-C, Supplementary Table 3). Gene ontology enrichment analysis of the differentially expressed genes identified cell type-specific processes relevant to pulmonary fibrosis (Figure 3D-F). To determine whether integrated single-cell RNA-seq analysis can recapitulate gene expression signatures previously identified in patients with pulmonary fibrosis, we used Gene Set Enrichment Analysis to compare the differential gene expression signatures we observed in alveolar macrophages, alveolar type II cells and fibroblasts identified from the integrated object, with genes associated with pulmonary fibrosis in the Comparative Toxicogenomics Database Pulmonary Fibrosis Gene Set (*17–19*). All three cell populations showed significant enrichment for genes in this dataset with normalized enrichment scores of 1.57, 2.02, and –1.79 for macrophages, alveolar type II cells and fibroblasts, respectively (FDR q-value < 0.01) (Figure 3G-L). Moreover, all examined cellular populations demonstrated changes in gene expression associated with pulmonary fibrosis (Supplementary Figure S3).

**Figure 3.**
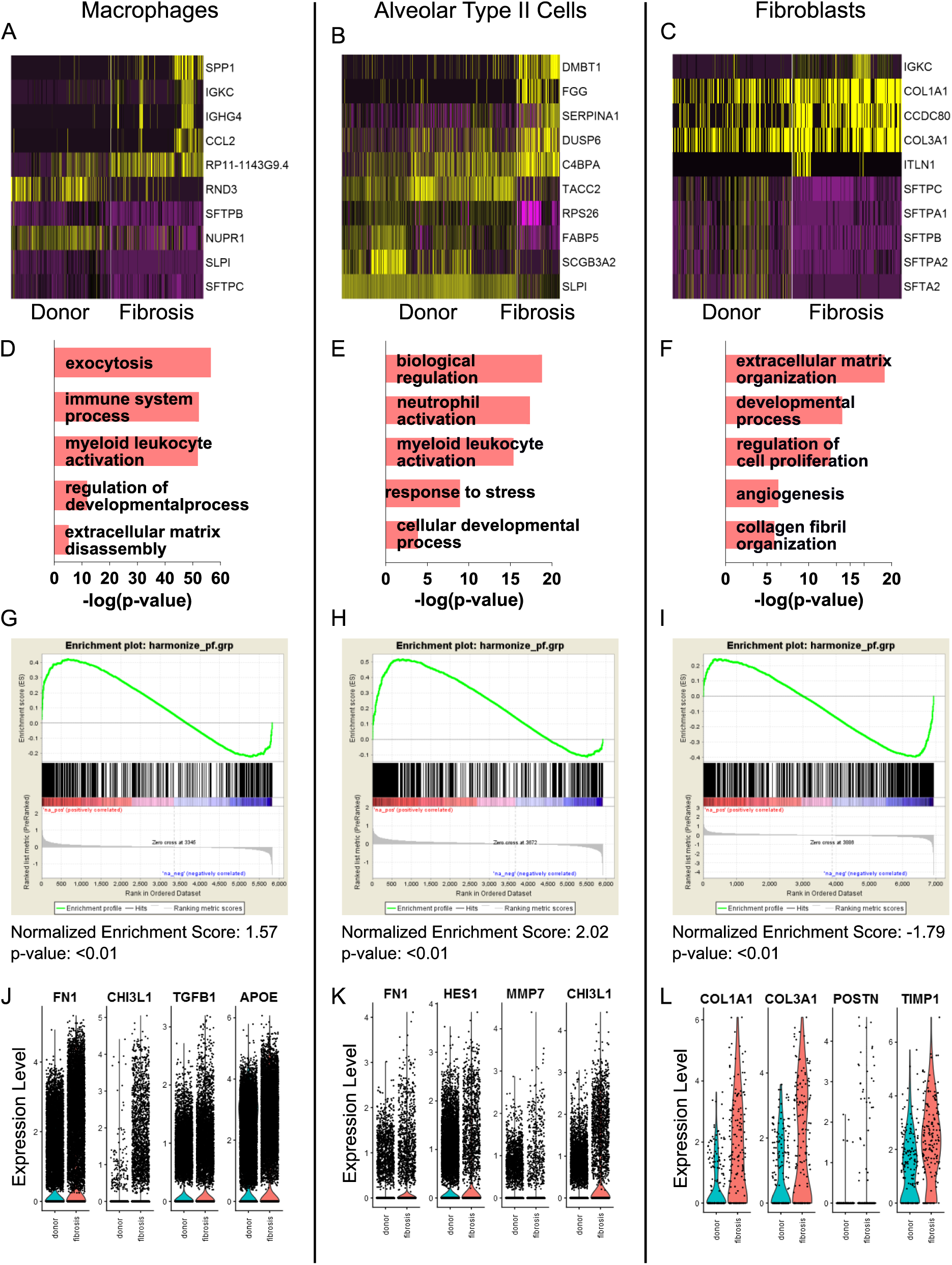
Differential expression analysis comparing donors and patients with pulmonary fibrosis identifies genes characteristic of pulmonary fibrosis. A-C, Differential expression analysis using Seurat’s FindMarkers function (Wilcoxon rank sum test) was performed comparing cells from donors and patients with pulmonary fibrosis within each cell type.

Heatmaps are shown representing the top five up- and down-regulated genes in macrophages, alveolar type II cells, and fibroblasts. D, E, Functional enrichment analysis with GO Biological Processes was performed using GOrilla with the top 1,000 genes upregulated in pulmonary fibrosis compared with donors. Representative significantly enriched GO processes are shown for macrophages, alveolar type II cells, and fibroblasts. G, H. Gene Set Enrichment Analysis was performed for the Comparative Toxicogenomics Database Pulmonary Fibrosis Gene Set using genes ranked by log difference in average expression between pulmonary fibrosis and donor subjects. Enrichment plots together with normalized enrichment scores and FDR q-values are shown for macrophages, alveolar type II cells, and fibroblasts. J-L. Violin plots of expression for select genes significantly upregulated in patients with pulmonary fibrosis compared with donors.

### Localization of known pulmonary fibrosis-associated signaling pathways to specific cell populations

The pathobiology of pulmonary fibrosis in animal models involves the aberrant activation of developmental pathways in the lung, including Notch, Wnt/β-catenin, and signaling pathways associated with epithelial to mesenchymal transition (EMT) (*20*). Our single-cell RNA-seq data allowed us to localize the expression of genes associated with these pathways to specific cellular populations within the diseased lung. Using donor lung gene expression as a control, we examined changes in the expression of genes in these pathways among epithelial and mesenchymal cell populations and macrophages as represented in dot plots and violin plots for selected genes (Figures 4A-E). Importantly, we were able to detect these genes in all subjects.

**Figure 4.**
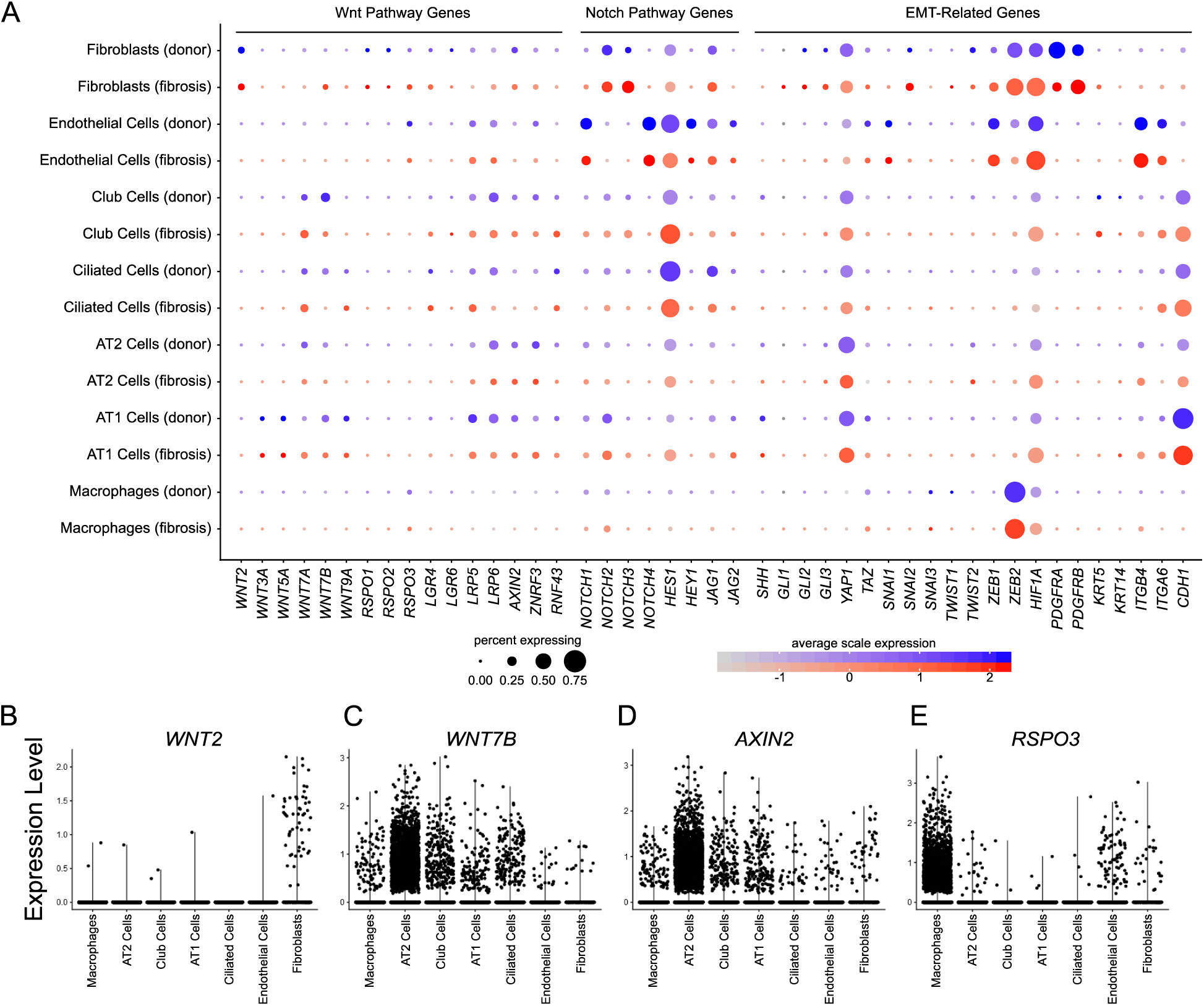
Single-cell transcriptomic analysis reveals distinct contributions of individual cell populations to pathways implicated in the pathogenesis of pulmonary fibrosis. A. Expression of selected Wnt pathway, Notch pathway, and epithelial to mesenchymal transition (EMT)-related genes is shown in fibroblasts, endothelial cells, club cells, ciliated airway cells, alveolar type II cells, alveolar type I cells and macrophages, separated by donor (blue) or pulmonary fibrosis (red) origin. Dot size corresponds to the percentage of cells in the cluster expressing a gene, and dot color corresponds to the average expression level for the gene in the cluster. B. Violin plots of *WNT2*, *WNT7B*, *AXIN2* and *RSPO3* expression are shown, suggesting a set of Wnt-expressor and -responder cell populations in the human lung.

We saw upregulation of Notch ligands and Notch target gene expression in alveolar type II cells and club cells, with downregulation of Notch target gene expression in endothelial cells. In this larger dataset, we observed expression of several Wnt ligands in epithelial cell populations in both the donor and fibrotic lung, as well as fibroblasts in both the human and mouse lung, which were not observed in a recent smaller single-cell RNA-seq dataset from mice (Supplementary Figure S4) (*21*). The Wnt target gene *AXIN2* was expressed in a minority of epithelial cells in the donor lung, and showed only modest increases during fibrosis. Several genes related to EMT were upregulated in fibroblasts during fibrosis. R-spondins are structurally distinct from Wnts, but work synergistically to sustain Wnt/β-catenin signaling (*22*). As expected, we found expression of R-spondins in fibroblasts, endothelial cells and surprisingly, in macrophages. None of the populations expressing R-spondins express Wnt ligands or the Wnt target, *AXIN2*.

These findings are consistent with the emerging concept that Wnt signaling, particularly in progenitor populations, is optimized through defined epithelial/mesenchymal pairings that comprise the niche (*21, 23, 24*). Accordingly, we examined our dataset for cells responsible for maintaining the progenitor niche. In the gut, there is a Wnt-responsive epithelial progenitor population defined by the R-spondin receptor *LGR5*, and this receptor along with *LGR6* has recently been reported to mark a mesenchymal population in the lung necessary for maintenance of a progenitor niche (*25, 26*). While we did not detect *LGR5* in the normal or fibrotic human lung or in the naïve adult mouse lung, possibly due to underrepresentation of mesenchymal cells, we detected *LGR4* and *LGR6* in a small subset of epithelial cells and fibroblast/mesenchymal cells (Figure 4A, Supplementary Figure S4). In mice, a subset of lung fibroblasts expressing *Pdgfra*, *Porcn*, *Wls*, *Wnt5a* and other Wnts was reported to form a niche responsible for maintenance of a rare population of *Axin2* positive alveolar type II cells that differentiate into type I cells during injury (*21*). These genes were detectable in fibroblasts from human and murine lungs (Figure 4A, Supplementary Figure S4).

To validate the single-cell RNA-seq analysis, we performed bulk RNA-seq analysis of flow-sorted alveolar macrophages, alveolar type II cells and whole lung tissue in a separate cohort of 14 donors compared with 8 patients with pulmonary fibrosis (Tables 1 and 2). Estimation of differential gene expression identified 3,817 genes in whole lung tissue, 4,891 in alveolar type II cells, and 1,238 in alveolar macrophages (Figure 5A-C). Hierarchical clustering of samples and identification of differentially expressed genes between donors and subjects with pulmonary fibrosis was performed (Figure 5D-F). Consistent with the single-cell RNA-seq data, whole lung tissue, alveolar macrophages and alveolar type II cells from patients with pulmonary fibrosis were significantly enriched for genes in the curated Comparative Toxicogenomics Database Pulmonary Fibrosis Gene Set (Figure 5J-O).

**Figure 5.**
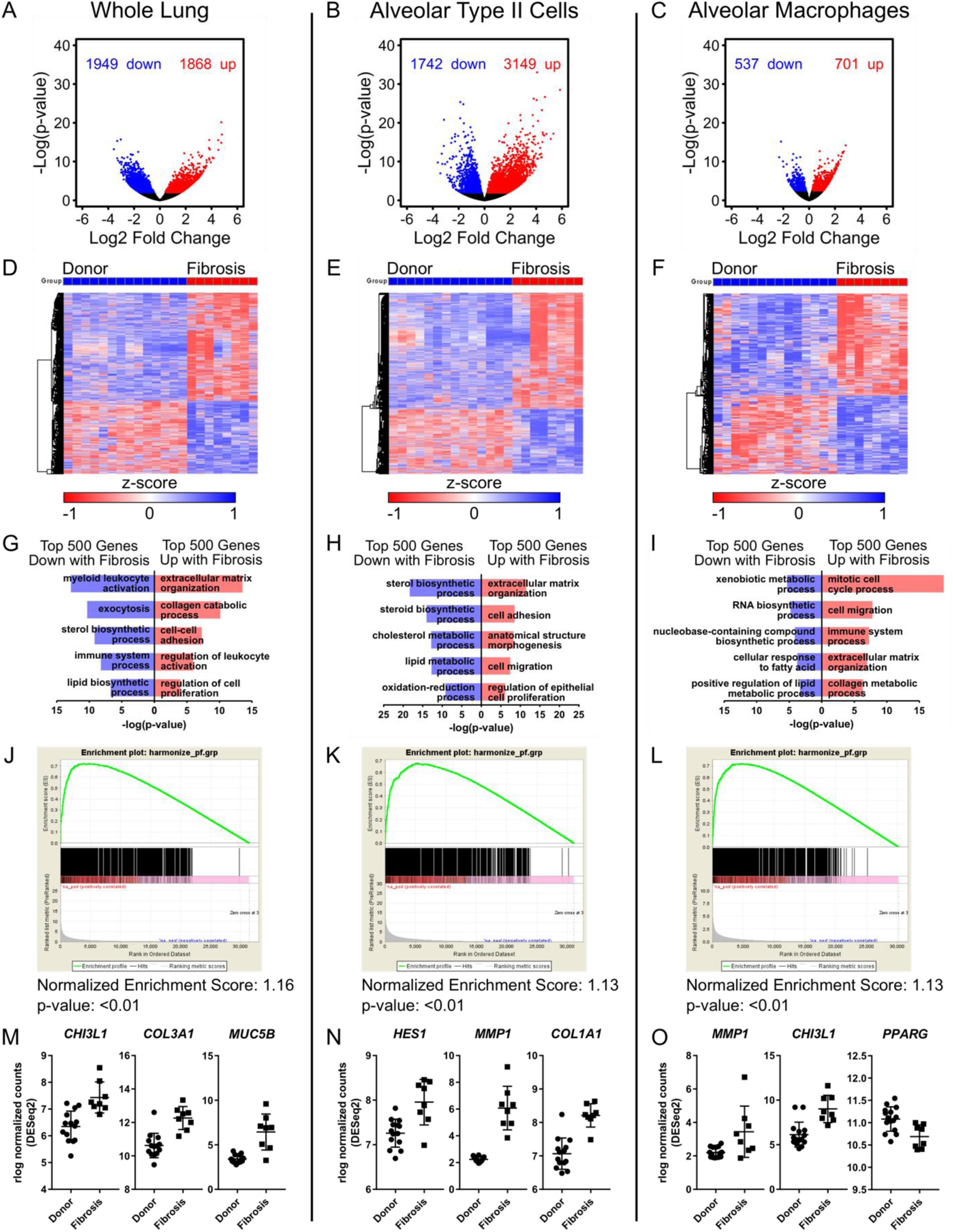
Transcriptional profiling of flow-sorted alveolar type II cells and alveolar macrophages from donors and recipients with pulmonary fibrosis validate the single-cell analysis. A-C. Lung tissue was obtained from 14 donors and 8 patients with pulmonary fibrosis at the time of lung transplantation. Single-cell suspensions were prepared and alveolar type II cells and alveolar macrophages were isolated using fluorescence-activated cell sorting, and transcriptional profiling using RNA-seq was performed on whole lung, alveolar type II cells, and alveolar macrophages. Estimation of differential gene expression using DESeq2 was performed comparing donors and patients with pulmonary fibrosis. Volcano plots are shown for whole lung, alveolar type II cells, and alveolar macrophages, respectively. D-F. Hierarchical clustering heatmaps of significant differentially expressed genes were generated using GENE-E. G-I. Functional enrichment analysis with GO Biological Processes was performed using GOrilla with the top 500 genes up-and downregulated in subjects with pulmonary fibrosis versus donor subjects. Representative GO processes are shown. J-L. Gene Set Enrichment Analysis was performed for the Comparative Toxicogenomics Database Pulmonary Fibrosis Gene Set using genes ranked by –log(FDR q-value). Enrichment plots together with normalized enrichment scores and FDR q-values are shown for whole lung, alveolar type II cells and alveolar macrophages. M-O. Expression of selected significant differentially expressed genes previously described to be important in pulmonary fibrosis.

### Single-cell RNA-seq data from individual patients identifies heterogeneity during pulmonary fibrosis

As an alternative approach to our integrated analysis of single-cell RNA-seq data using CCA, we separately identified cell clusters in each of the individual subjects. This approach allowed us to include all 8 donors and 8 patients with pulmonary fibrosis (Supplementary Figure S5 contains individual t-SNE plots for each patient; Supplementary Table 4 includes the marker genes used for cell assignment). Analysis of the individual subjects revealed several distinct populations that were not observed in the integrated analysis. In total, we identified 22 distinct cell types by individual annotation of each subject, compared with 13 cell populations identified by the integrated analysis. Many of the populations not observed using the integrated analysis were represented only in a minority of patients or in a single patient, and were combined with other cell populations in the integrated analysis. For example, in a patient with pulmonary fibrosis (Supplementary Figure S6A, Subject IPF 2), we identified a cluster of *FOXJ1*-negative epithelial cells in which the most highly expressed genes included *SFTPC*, *SFTPB*, *SFTPA2* and *LAMP3*, suggesting these cells are alveolar type II cells. Nevertheless, these same cells also expressed the club cell markers *SCGB3A2*, *SCGB3A1*, and *SCGB1A1*. These cells also expressed high levels of *HES1*, *MMP7* and other genes associated with fibrosis. Furthermore, in the same subject we could resolve *FOXP3* and *CD4* expressing regulatory T cells, and *ITGAE* and *CD8* expressing lung resident T cell subsets. Similarly, in the patient with hypersensitivity pneumonitis, we identified two small populations of CD45 expressing cells that were not detected in any other subject (Supplementary Figure S6B). The first cluster (30 cells) expressed *CLEC4C*, *TCL1A*, *IRF8* and *TLR7*, markers associated with plasmacytoid dendritic cells. The second cluster (31 cells) expressed *CLEC9A*, a typical marker of a subset of conventional dendritic cells (DC1).

These populations were classified as macrophages in the integrated analysis. The distinction between classical and non-classical monocytes, which was apparent in the subjects IPF 1, HP, Donor 6 and SSc-ILD 1 were not identified in the integrated analysis (Supplementary Figure S6C). It is therefore possible the improved resolution in the individually annotated analysis is driven by changes in cellular composition between donor and fibrotic lungs, the emergence of new cell types in patients with fibrosis, and relative under-sampling of some cell populations. We re-examined alveolar macrophages and alveolar epithelial cells by combining cells identified by individual annotation from each patient. To capture potentially novel epithelial cell populations in patients with fibrosis, we included all epithelial populations (airway and distal lung) in our analysis.

### Alveolar macrophages

In a murine model of lung fibrosis induced by the intratracheal administration of bleomycin, we recently reported that alveolar macrophages during fibrosis are comprised of two ontologically distinct populations—tissue-resident alveolar macrophages and monocyte-derived alveolar macrophages (*27*). In that system, we showed that genetic deletion of monocyte-derived, but not tissue-resident, alveolar macrophages improved fibrosis (*27*). Bulk RNA-seq analysis identified a fibrotic gene expression signature in the monocyte-derived cells, a finding that has been subsequently confirmed by an independent group (*28*). These studies predict the presence of homeostatic tissue-resident alveolar macrophages in the normal lung, and both tissue-resident alveolar macrophages and profibrotic, monocyte-derived alveolar macrophages in the fibrotic lung. We combined clusters of cells identified as macrophages from each of the 8 donors and 8 patients with fibrosis and performed unbiased clustering of these cells. After removal of contaminating cells, clustering was performed and four distinct macrophage clusters were identified (Figure 6A, Supplementary Table S5). Interestingly, Cluster 0 contained almost all of the alveolar macrophages from the donor lungs, but only a fraction of alveolar macrophages from patients with pulmonary fibrosis, irrespective of the diagnosis (Figure 6B, C). In contrast, Clusters 1 and 2 originated largely from the lungs of patients with fibrosis (Figure 6C). Cells in Cluster 1 expressed lower levels of *PPARG, MRC1* and *MARCO,* and higher levels of *APOE* and *MAFB*, which we and others have reported rise and fall, respectively, as monocytes differentiate into alveolar macrophages (Figure 6D) (*27, 29*). To explore these clusters further, we compared genes differentially expressed in each cluster with the average gene expression in all macrophages using functional enrichment with GO Biological Processes. Cluster 1 included processes associated with fibrosis including “exocytosis”, “secretion”, “regulation of cell migration” and “extracellular matrix organization”. In contrast, upregulated processes in Cluster 0 were suggestive of homeostatic functions of macrophages including “immune system processes”, “response to lipid” and “response to organic/inorganic substance”. Processes upregulated in Cluster 2 overlapped with those in Cluster 1, but did not include cell migration or matrix reorganization. Finally, processes in Cluster 3 were implicated in the metabolism of metals. A full list of these GO processes is provided in Supplementary Table 5.

These single-cell RNA-seq data suggested heterogeneity in the expression of profibrotic genes in alveolar macrophages from the same patient. We tested this hypothesis by performing immunohistochemistry on lung sections from tissue adjacent to the biopsy processed for single cell RNA-seq at the time of harvest. We selected *CHI3L1*, *MARCKS*, *IL1RN*, *PLA2G7*, *MMP8* and *SPP1,* as they are differentially expressed in alveolar macrophages between subjects with pulmonary fibrosis and donors, are associated with pulmonary fibrosis in the Comparative Toxicogenomics Database Pulmonary Fibrosis Gene Set, and have validated antibodies for use in the Human Protein Atlas (*30*). Consistent with the single-cell RNA-seq data, we found that alveolar macrophages from donor lungs did not stain for these markers, and only a subpopulation of cells in any given patient with pulmonary fibrosis expressed profibrotic proteins (Figure 6E). Although our sampling protocols are insufficient to make firm conclusions, the relative expression of a given marker in a single patient on RNA-seq (abundant or infrequent), was reflected in the immunohistochemistry. Moreover, even with a small number of patients in our cohort, we identified that expression of the individual profibrotic markers in alveolar macrophages, was highly variable among the patients with the same clinical diagnosis (Figure 6F). This finding is important as alveolar macrophages can be sampled safely and repetitively over the course of pulmonary fibrosis through bronchoscopic lavage.

### Epithelial Cells

Epithelial cells contained clusters representing the major known lung epithelial cell types: ciliated epithelial cells (*FOXJ1)*, club cells (*SCGB1A1*), alveolar type I cells (*AGER*) and alveolar type II cells (*SFTPC*, *LAMP3*). Ciliated cells and alveolar type I cells isolated from patients with pulmonary fibrosis clustered well with cells isolated from donors (Figure 7A-D, Cluster 2 and Cluster 5). As we observed with alveolar macrophages, cells expressing canonical alveolar type II cell markers from donors formed a single cluster that also included cells from patients with pulmonary fibrosis (Figure 7A-D, Cluster 0). In contrast, it was difficult to use canonical markers to clearly identify unique epithelial cell populations that emerged in patients with pulmonary fibrosis (Figure 7A-D, Clusters 1 and 2). While some of these cells expressed moderate or high levels of *SFTPC*, these cells also expressed genes typically associated with club cells, suggesting the emergence of intermediate epithelial cell phenotypes during pulmonary fibrosis. One of these clusters (Cluster 2) is particularly illustrative in that it includes cells from most of the patients with pulmonary fibrosis and is characterized by high levels of expression of *SFTPC*. We generated violin plots using common genes from Cluster 0, with *CHI3L1* and *SFTPC* shown as examples of genes with relatively high and low heterogeneity, respectively (Figure 7E).

**Figure 7.**
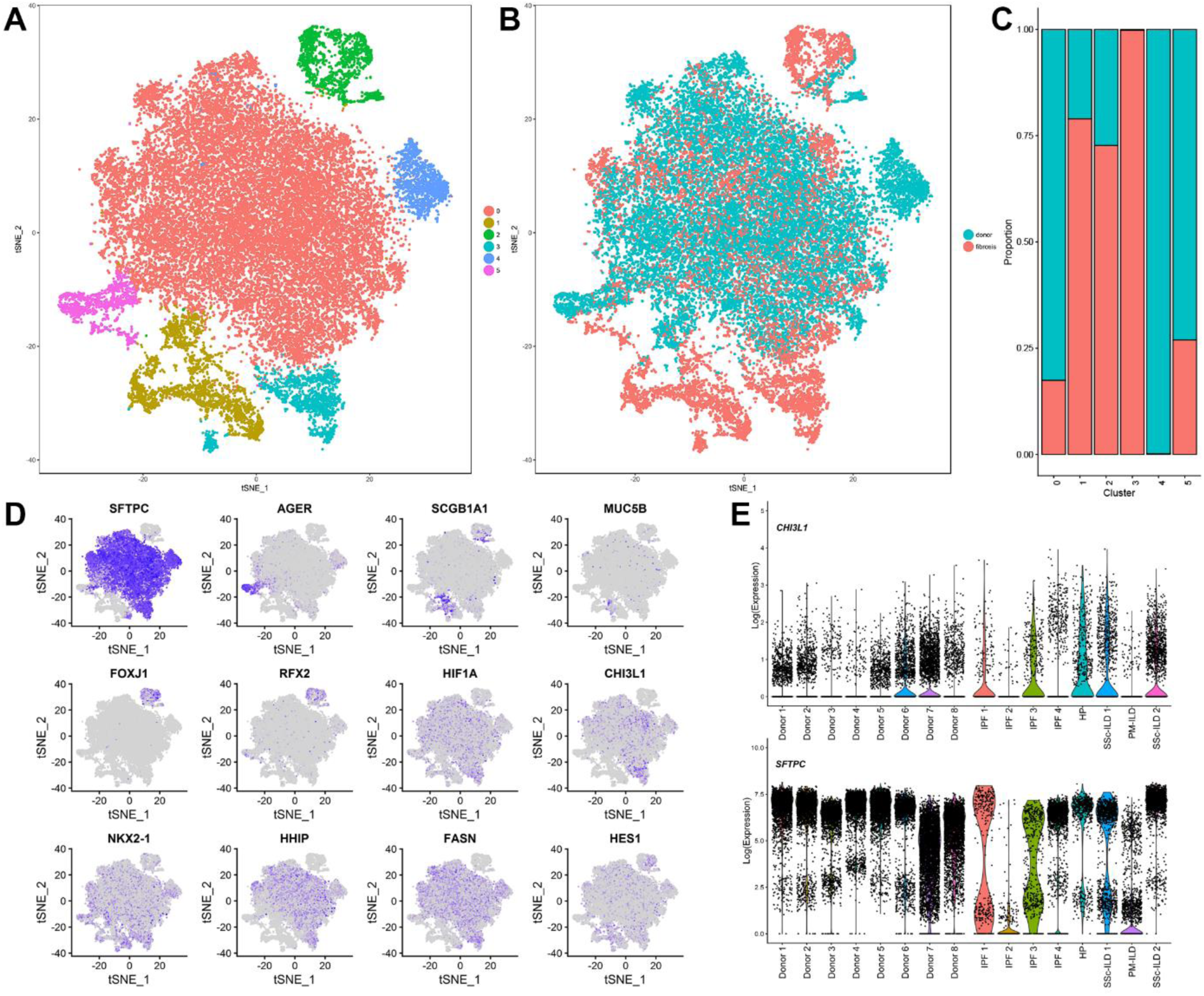
Distinct populations of alveolar epithelial cells emerge during fibrosis. A. Six clusters were identified after epithelial cells from each of the 16 individual subjects (8 donor and 8 fibrosis) were combined and clustered. B. t-SNE plot demonstrating the contribution of cells from the lungs of donors and patients with fibrosis to each cluster. C. Relative contributions epithelial cells form the lungs of donor and patients with fibrosis to each cluster. D. Feature plots demonstrating differential expression of selected epithelial marker genes: *SFTPC* (alveolar type II cells), *AGER* (alveolar type I cells), *SCGB1A1* (club cells), *FOXJ1*, *RFX2* (ciliated airway epithelial cells). Also shown are genes implicated in pulmonary fibrosis (*HIF1A*, *CHI3L1*, *NKX2-1*, *HHIP*, *FASN*, *HES1*). E. Violin plots demonstrate heterogeneity in gene expression of *CHI3L1* and *SFTPC* in epithelial cells from the lungs of donors and patients with fibrosis.

Since our data show significant heterogeneity in epithelial cells between both donors and patients with lung fibrosis, we re-examined expression of Wnt/β-catenin pathway components during fibrosis in the combined epithelial cluster in more detail. Likely due to increased sampling, we were able to identify higher rates of *AXIN2* expression in alveolar type II cells in both mice and humans than were recently reported (*21*) (Supplementary Figures S4, S7). We found low levels of expression of most Wnt ligands in epithelial cells from humans, but *WNT5A* and *WNT7B* were by far the most abundantly expressed (Supplementary Figure S7). Among queried Wnt ligands, we found non-overlapping patterns of expression between cell populations that express Wnt ligands and the canonical Wnt target, *AXIN2* (Figure 8A, B). Furthermore, while many cells within the lung express Wnt ligands, including alveolar type II cells, individual cells to a large extent express a single Wnt ligand (Figure 8C). We confirmed these findings using *in situ* RNA hybridization in human tissues. Consistent with the single-cell RNA-seq data, airway epithelia largely expressed *WNT7B*, but we did not detect expression of *AXIN2*. In the alveolar epithelium, we confirmed increased expression of *WNT7B* relative to *WNT7A*, and found that cells expressing either *WNT7A* or *WNT7B* were generally distinct from those expressing *AXIN2* (Figure 8D-G). This distinction was not absolute, as we were able to identify a small number of alveolar epithelial cells expressing both *WNT7B* and *AXIN2* (Figure 8G).

**Figure 8.**
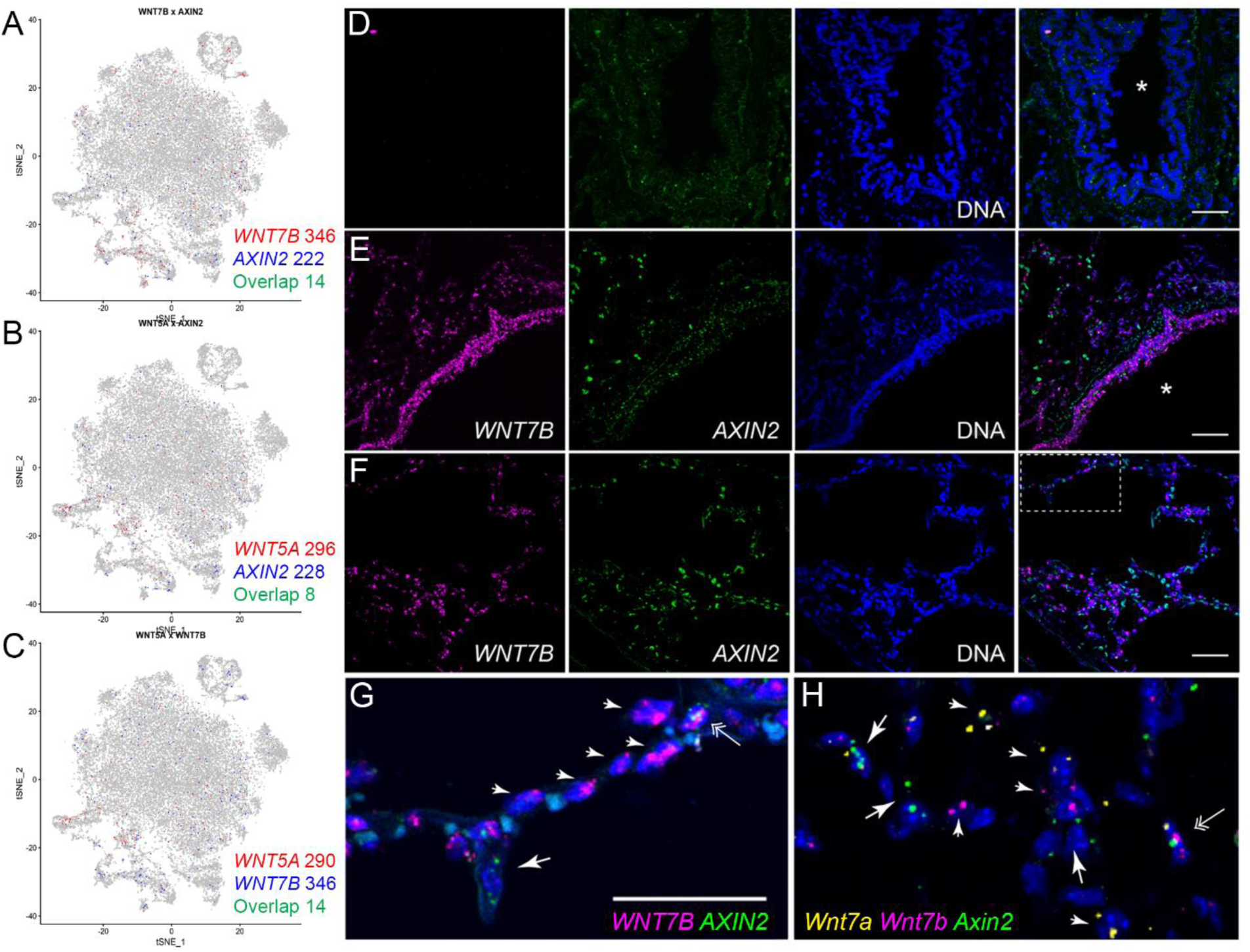
Wnt secretion and response are restricted to distinct non-overlapping cells. A-C. Expression of *WNT7B, WNT5A* and *AXIN2* in epithelial cells from lung of donors and patients with pulmonary fibrosis by single-cell RNA-seq. D, E. Fluorescence RNA *in situ* hybridization reveals Wnt-expresser and -responder cells in the proximal airways of human lung. Images were obtained from human donor lung sections incubated without (D) or with designated target probes, *WNT7B* (magenta) and *AXIN2* (green) (E-G) Nuclei are stained with Hoechst (blue) and overlay images are shown on the far right. Note *WNT7B* is abundantly detected in small airway epithelial cells (E), consistent with its prominent detection in club cells by single-cell RNA-seq. *AXIN2* is not similarly enriched in club cells, and is only specifically detected in alveolar regions (F) under higher magnification (G is a higher magnification image of the boxed region in F). H. High magnification of mouse lung alveoli subjected to fluorescence RNA *in situ* hybridization with probes for *Wnt7a* (yellow), *Wnt7b* (magenta) and *Axin2* (green). Arrows indicate cells that are only positive for *Axin2*; arrowheads indicate specified Wnts. Double-arrows indicate cells that may express both Wnts and *Axin2*, although quantification suggests cells that highly express Wnts and Axin2 are rare (Supplementary Figure S8). Scale bar 50 μm. Asterisk (*) denotes airway lumen.

The observation that Wnt secretion and response are largely restricted to distinct non-overlapping cells in the human lung was confirmed by single-cell RNA-seq analysis of mouse lung. Specifically, murine alveolar type II cells expressed *Wnt3a*, *Wnt7b*, *Wnt4* and *Wnt9a*, whereas alveolar type I cells expressed *Wnt3a*, *Wnt7a*, *Wnt9a* and *Wnt10b* (Supplementary Figure S4). As in the human lung, we mostly detected only one Wnt-ligand at a time (Supplementary Figure S8A). We also confirmed that Wnt-ligand and *Axin2* expression in mouse alveolar type II cells appear largely mutually exclusive by both single-cell RNA-seq (Supplementary Figure S8B) and *in situ* RNA hybridization analysis (Figure 8H, Supplementary Figure S8C, D).

### Single-cell RNA-seq identifies rare cell populations in human lung

We queried our data for evidence of gene expression associated with stem cell populations and senescence. In the airway, investigators identified a population of basal cells in the airway that expand after tracheal injury (*31*). Using transcriptional profiling, we identified a cluster of cells expressing many of the genes distinctly associated with these cells including *KRT5*, *TP63*, *SCGB1A1* and *SOX2* with variable expression of *ITGA6* and *NGFR* (Figure 9A). Two recent reports identified a sub-population of alveolar type II cells expressing high levels of Axin2 (2-20% of mouse and 29% of human alveolar type II cells), and demonstrated these cells serve as a facultative progenitor cell population that repopulates the lung after injury (*21, 32*). Although Axin2-positive alveolar type II cells express all of the established markers of this cell type (*21, 32*), the study by Zacharias et al. identified 875 and 2,773 genes that were differentially expressed in the *AXIN2* positive progenitor cell population relative to *AXIN2* negative alveolar type II cells in the human and mouse lung, respectively. Since single-cell RNA-seq is able to identify much smaller differences in gene expression between populations, we searched for these populations in the normal mouse alveolar type II cells. However, unbiased clustering of the 1,821 alveolar type II cells we sampled did not identify subclusters of alveolar type II cells enriched for expression of *Axin2* or *Tm4sf1* (Supplementary Figure SA-D). To provide additional power to this guided analysis, we used the 500 reported differentially expressed genes which were reported to be upregulated in *Axin2*-positive progenitor cells as a gene module for the AddModuleScore function implemented in Seurat package to find the cells enriched for these genes. We did not find subclusters of alveolar type II cells enriched for the genes associated with *Axin2*-positive progenitors in the previous studies (Figure 9E-F). We then searched for this sub-population within alveolar type II cells from each of the eight donor lungs and performed unbiased clustering, which did not reveal additional sub-clusters (Supplementary Figure S9G). These data suggest that single-cell RNA-seq can readily identify transcriptional differences between dedicated progenitor populations (e.g. basal cells), and that alveolar type II cells in normal mouse and human lung represent a relatively homogenous population.

**Figure 9.**
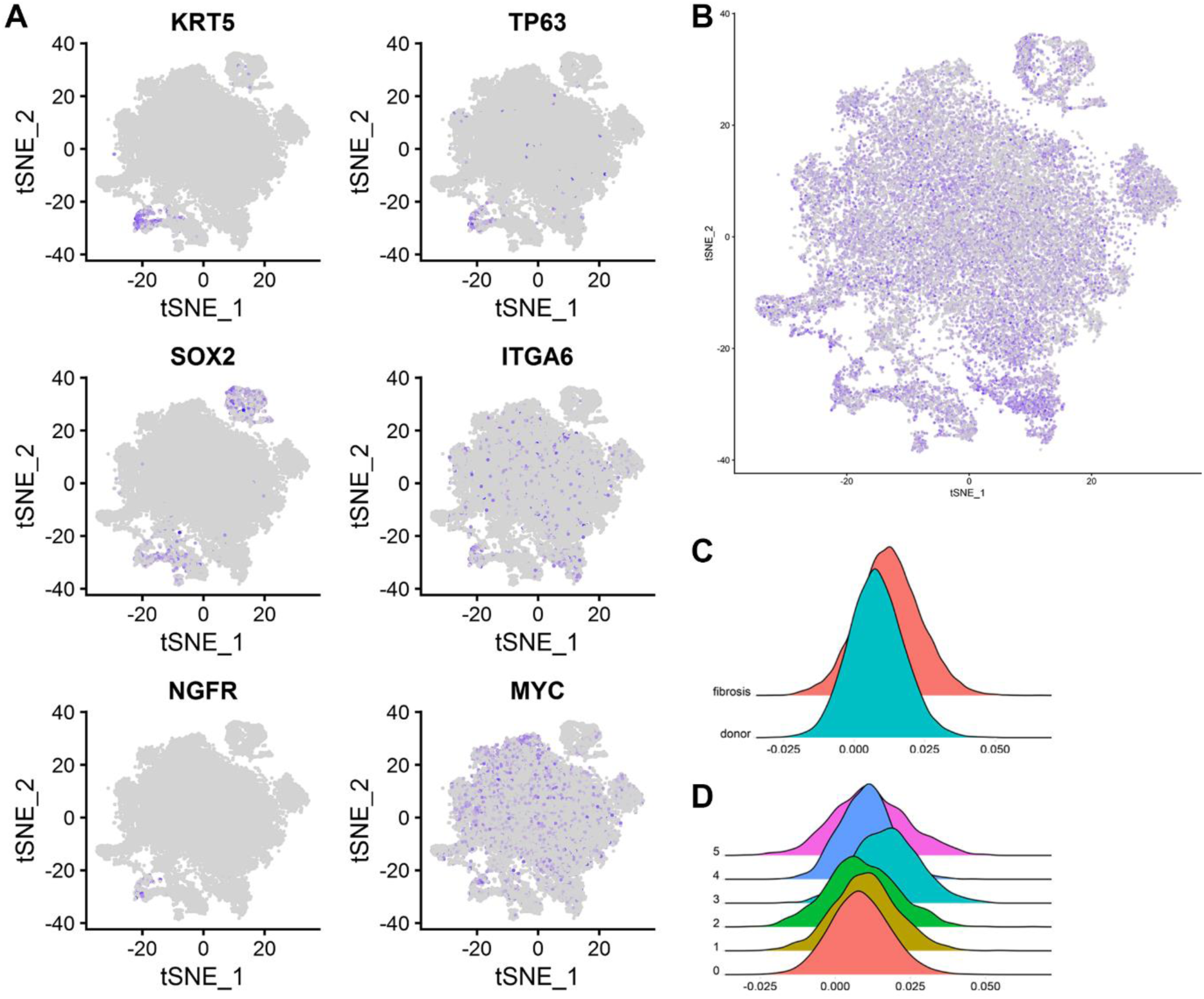
Identification of rare cell populations among lung epithelial cells using single-cell RNA-seq. A. Expression of markers associated with airway basal stem cells. B. Feature plot showing cells enriched for senescence-associated genes using a senescence score for each cell.

Terminally differentiated cells in culture undergo a relatively constant number of population doublings before undergoing replicative senescence. The development of senescence is associated with characteristic changes in gene expression that include the induction of p16, p21 and p53, which slow the cell cycle and promote resistance to apoptosis. Senescence also results in the production and secretion of a distinct set of proteins collectively referred to as senescence-associated secretory proteins (*33*). These include insulin-like growth factor-binding proteins, interleukins, transforming growth factor type-β and plasminogen activator inhibitor-1. A growing body of evidence supports a link between aging, cellular senescence in the lung and the susceptibility to lung disease. For example, investigators have reported increased markers of senescence in the lungs of patients with IPF, and have found that myofibroblasts obtained from the lungs of patients with IPF demonstrate a senescence-associated resistance to apoptosis (*34, 35*). However, even in pulmonary fibrosis, the percentage of cells in the lung that are positive for senescence markers is very low, precluding their identification using bulk RNA or protein analysis. Hernandez-Segura and colleagues have performed a meta-analysis of the transcriptional changes associated with replicative senescence in culture (*36*). We used this set of 1,311 genes to generate a senescence score for any individual cell and applied it to epithelial cells in our dataset. We observed increased average expression for senescence-associated genes in the population of alveolar type II cells uniquely present in patients with pulmonary fibrosis (Figure 9B). We used the average expression of these genes to generate a senescence score for each of the epithelial populations in our combined analysis (see Figure 7 for a description of the Clusters). We then generated a combined senescence score for all epithelial cells for each patient with pulmonary fibrosis and each donor and found this score was significantly higher in patients with pulmonary fibrosis (Figure 10C, p = 0.0001). The senescence score was highest in the unique population of alveolar type II cells in patients with fibrosis (Cluster 3, Figure 9D). These results suggest that after validation with lineage tracing studies in mice, single-cell RNA-seq offers promise for the quantification of senescent cells in the lung.

**Figure 6.**
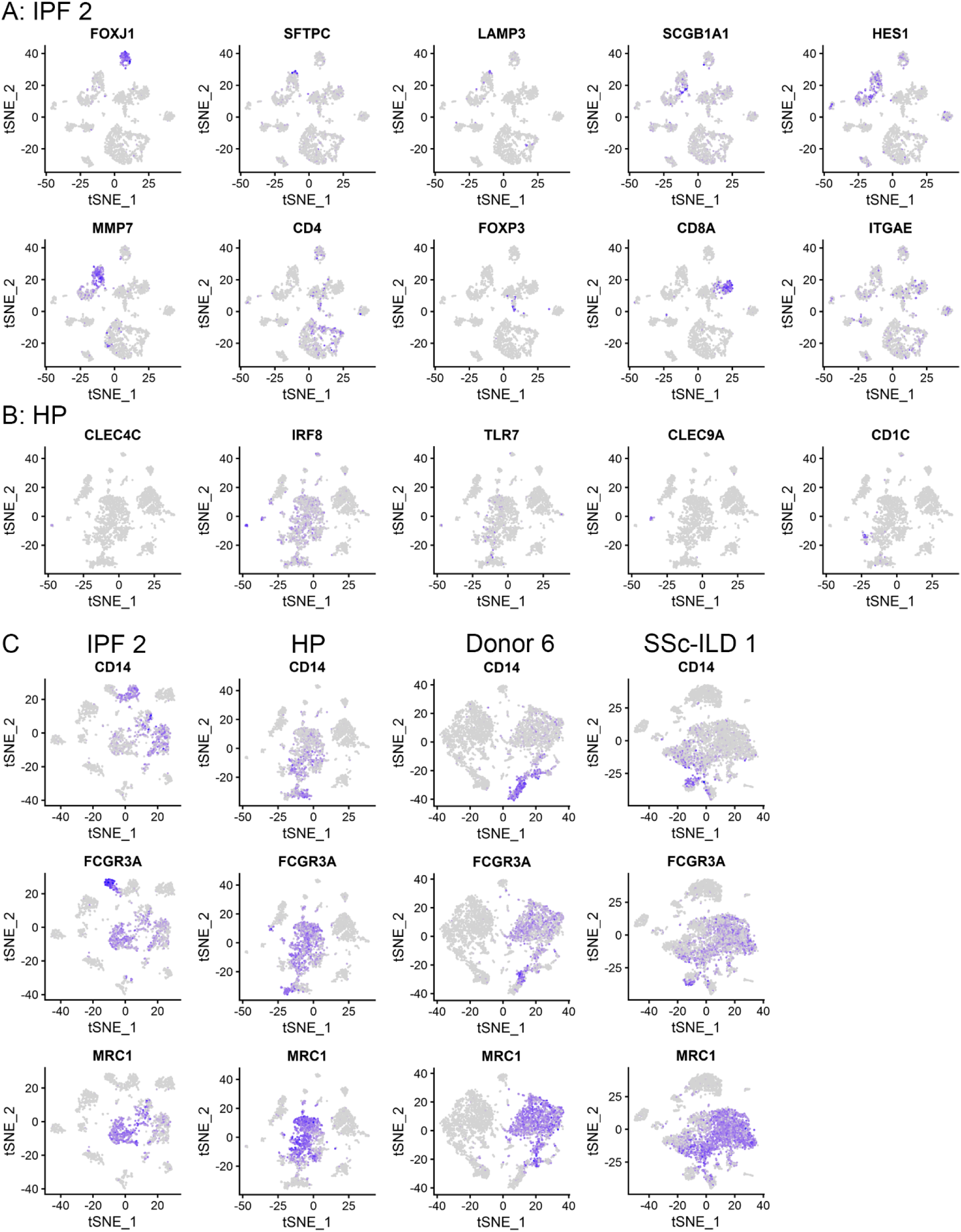
Distinct populations of alveolar macrophages emerge during lung fibrosis. A. Individually annotated alveolar macrophage clusters from each of the 16 individual subjects (8 donors and 8 fibrosis) were combined to create a single object and then clustered, revealing four clusters. B, C. Relative contributions of alveolar macrophages from the lungs of donors and patients with pulmonary fibrosis to each cluster as shown by t-SNE plot and by bar plots. D. Feature plots demonstrating differential expression of selected alveolar macrophage maturation genes (*PPARG*, *APOE*, *MARCO*, *MRC1*, *MAFB* and *SIGLEC1)*, and genes associated with fibrosis (*IL1RN*, *MMP9*, *CHI3L1*, *SPP1*, *MARCKS*, *PLA2G7*). E. Immunohistochemistry on lung sections from the same patients confirms heterogeneity in fibrotic gene expression within alveolar macrophages from the same patient. Arrows indicate positive staining in alveolar macrophages, arrowheads indicate positive staining in alveolar epithelium for *CHI3L1* and *SPP1*, endothelium for *MARCKS* and neutrophils for *MMP9*. F. Heterogeneity of fibrotic gene expression in alveolar macrophages between subjects.

C. Histograms showing the average senescence-associated gene expression score in all epithelial cells from patients with lung fibrosis compared with donors (P=0.0001, t-test, n=8 donor and fibrosis). D. Histograms showing average senescence score by Cluster (see Figure 7 for cluster details).

### Single-cell RNA-seq of lung biopsy tissue from a living patient

We next examined the feasibility of single-cell analysis from a cryobiopsy specimen obtained *ex vivo* from an explanted lung specimen from a patient with IPF using flow cytometry. We were able to resolve alveolar macrophages and alveolar epithelial cells, both with good viability (Supplementary Figure S10). Accordingly, we obtained a cryobiopsy specimen from a patient at the time of video-assisted thoracoscopic biopsy. The patient was subsequently diagnosed with IPF based on histopathological examination of the biopsy (Figure 10A,B). We identified 1,516 cells corresponding to 13 different cell populations (Figure 10C). Among the detected cells, we were able to resolve a distinct population of cells characterized by expression of *PROX1, MMRN1, TBX1* and *RELN* genes, suggesting that these cells represent lymphatic progenitors, which were underrepresented in samples obtained at the time of transplant (Figure 10D).

**Figure 10.**
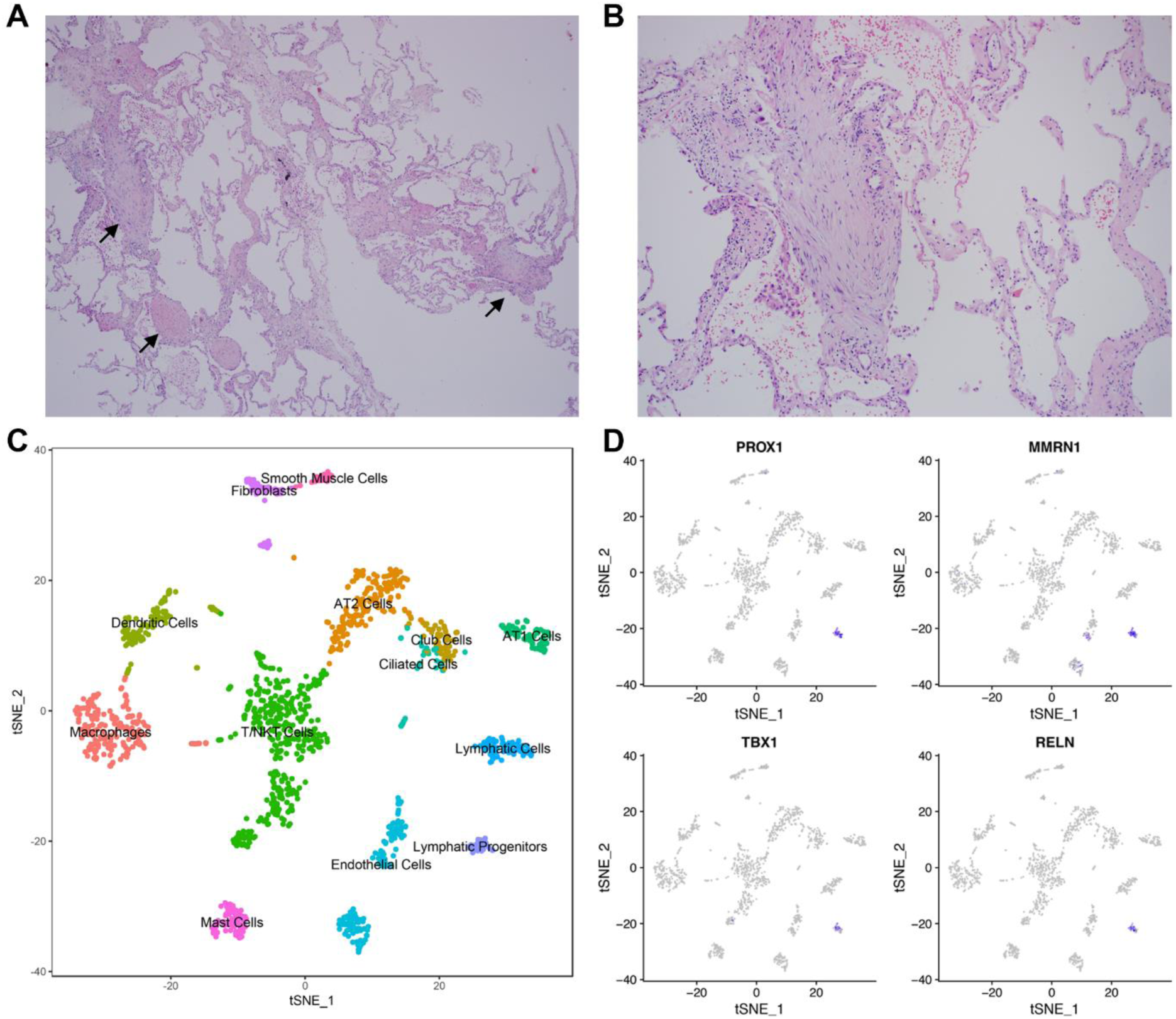
Single-cell RNA-seq of a cryobiopsy from a living patient with IPF. A, B. Representative histology (H&E) showing fibroblastic foci (arrows), 40x and 100x magnification correspondingly. C. tSNE plot clustering of 1,516 cells into 13 distinct cellular types. D. Lymphatic progenitors were identified by expression of *PROX1, TBX1, RELN* and *MMRN1*.

## Discussion

We used two methods to perform an integrated single-cell transcriptomic analysis of lung tissue obtained from lung transplant donors and patients with pulmonary fibrosis. We detected changes in gene expression associated with pulmonary fibrosis across all identified cell populations, and illustrated how single-cell transcriptomic data could be used to dissect multicellular aberrant signaling processes implicated in the development of pulmonary fibrosis. We showed that changes in gene expression associated with fibrosis were readily detected in flow cytometry-sorted alveolar macrophages from the lungs of patients with pulmonary fibrosis. We demonstrated that single-cell transcriptomic analysis of tissue obtained bronchoscopically via cryobiopsy was feasible. Taken together, our findings suggest that the application of single-cell RNA-seq analysis to alveolar macrophages and/or lung tissue obtained during routine care, can be used to develop a molecular approach to improve the diagnosis and therapeutic monitoring of patients with pulmonary fibrosis.

The use of single-cell transcriptomics to study pulmonary fibrosis pathogenesis overcomes many of the challenges inherent in previous microarray or RNA-seq studies (*9–11, 37, 38*).

Transcriptomic analysis of whole lung tissue is heavily influenced by changes in the cellular composition of the tissue that mask changes in gene expression within cell populations. We found that both bulk RNA-seq applied to flow-sorted cell populations and single-cell RNA-seq analysis could overcome this problem. Flow sorting inherently assumes a relatively constant level of surface marker expression during health and disease, however few markers are explicitly validated for identification of cellular populations during disease states. This problem is exacerbated when antibodies are used for which the gene encoding the antigen is not known (*39*).

We found that single-cell RNA-seq applied to whole lung tissue, when combined with anatomic studies using immunohistochemistry or *in situ* hybridization techniques, allows the transcriptome-level reconstruction of multicellular anatomical niches important for disease pathogenesis or repair. For example, we identified a distinct population of alveolar macrophages during fibrosis that we predicted from genetic lineage tracing studies during fibrosis in mice (*27*), and we identified a population of alveolar type II cells enriched for senescence markers during pulmonary fibrosis (*40*). We used single-cell RNA-seq data to predict a distinction between cells expressing Wnt ligands and the canonical Wnt target *AXIN2/Axin2* in the normal and fibrotic lung, which we confirmed using *in situ* RNA hybridization. In addition, our data suggest lung macrophages as an unexpected potential source of R-Spondins that might augment Wnt signaling during fibrosis.

We also leveraged our data to support the conclusions of recently published studies. We were able to identify markers of some mesenchymal cell populations suggested to play a role in Wnt/β-catenin signaling during lung regeneration (*26, 41*). However, we did not observe any wide-scale upregulation of Wnts in samples from patients with pulmonary fibrosis, nor expansion of Wnt/Axin2-double positive alveolar type II cells as observed in the hyperoxia model of murine lung injury (*21*). We identified a likely population of *TP63* and *KRT5* expressing airway progenitor cells (basal cells) in the normal and fibrotic lung based on their transcriptome, and we largely confirmed the observation that *Axin2*-positive alveolar type II cells are a minor subset of the murine alveolar type II cell population (*21, 42*). Nevertheless, we were unable to identify distinct transcriptional signatures in *AXIN2*-expressing compared with non-expressing cells, perhaps suggesting that the distinct transcriptional signatures observed in dedicated progenitor populations are not present in facultative progenitors.

The falling costs of transcriptomic profiling make it a promising clinical tool to identify biomarkers that predict disease severity or response to therapy. Ideally, the cellular material used for transcriptomic profiling should be safe and easy to collect so that it can be readily sampled before and after the initiation of therapy. While peripheral blood fulfils these criteria, transcriptomic analyses of peripheral blood mononuclear cells have not generated markers of disease outcome with sufficient predictive accuracy to be adopted into clinical practice (*43*). To determine the feasibility of transcriptomic profiling in tissue samples obtained during routine patient care, we performed single-cell RNA-seq analysis on tissue obtained from a bronchoscopic cryobiopsy – a novel minimally invasive biopsy technique that is seen as a promising alternative to thoracic surgery especially in high-risk patients (*44*). Single-cell analysis of alveolar macrophages is attractive from a clinical perspective, as large numbers of cells can be safely and repeatedly sampled using bronchoscopic alveolar lavage, even in patients who are ill. Our analysis of alveolar macrophages suggests that single-cell RNA-seq identifies a distinct population of alveolar macrophages with higher expression of profibrotic genes, and accurately reveals differences in fibrotic gene expression in macrophages from different patients. Single-cell profiling of lung biopsies and/or alveolar macrophages from a larger number of patients might therefore identify patient-specific markers that could inform a personalized approach to therapy.

There are limitations to our study that highlight the need for further work to optimize and expand single-cell RNA-seq datasets for pulmonary fibrosis, and perhaps other diseases. First, while our single-cell transcriptomic dataset is the largest in the literature to date, it includes only a small number of donors and patients with pulmonary fibrosis. Even in this small cohort, however, we were able to identify many of the same genes that we detected in flow-sorted cell populations from an independent cohort of patients, as well as many of the genes identified in the literature. Second, the lung is a complex tissue, consisting of at least 40 cell types tightly embedded into extracellular matrix, and both the cellular composition and matrix characteristics change during disease. Isolation of intact cell populations faithfully representing the composition of the lung tissue via current enzymatic digestion protocols is therefore not feasible. Alternative approaches, such as single nucleus sequencing will likely be necessary to determine the cellular composition of the lung in disease (*45, 46*). Third, we have conducted our study on late stage disease, but we show that studies using biopsies obtained earlier are feasible and might identify pathways distinctly active early in the course of fibrosis. Fourth, computational approaches for work with single-cell transcriptomic datasets are in their infancy. As such, they are heavily biased by the experimentalists’ familiarity with the biology of the systems being studied, are not suitable for unbiased identification of rare populations, and are not yet adapted to correct for batch effect associated with multiple samples or disease conditions. Our findings suggest that the generation of larger datasets from normal and diseased samples, coupled with spatial information from RNA *in situ* hybridization and immunohistochemistry will drive improvements in these tools, and expand the current atlas of the cellular types and states.

In conclusion, single-cell RNA sequencing of normal and fibrotic lung tissue reveals shared as well as distinct patterns of fibrotic gene expression in individual cell populations and emphasizes the importance of intracellular communication in disease pathobiology. Single-cell transcriptomic analysis offers promise for the discovery of multicellular pathways important for disease pathogenesis, and for the generation or exclusion of hypotheses regarding the function of distinct cell populations that emerge during disease. Transcriptomic profiling of single cells can be performed using bronchoscopically obtained samples, and can show changes characteristic of pulmonary fibrosis. These technologies may therefore be used clinically to identify patients most likely to benefit from targeted therapy and to monitor their response over the course of disease.

## Supplementary Methods

### Patients

All procedures using human tissues were approved by the Northwestern University Institutional Review Board (STU00056197, STU00201137, and STU00202458) and by the University of Chicago Institutional Review Board (NCT00515567 and NCT00470327) and received further approval from The United States Army Medical Research and Materiel Command`s Office of Research Protections, Human Research Protection Office. All patients provided written informed consent.

### Mice

All procedures using mice were approved by the Northwestern Institutional Animal Care and Use Committee. Twelve week old male C57Bl/6 mice obtained from Jackson Laboratories were used for these experiments.

### Tissue processing and cell sorting

Upon arrival to the laboratory, a representative piece of whole lung tissue was collected for histology followed by fixation in 4% PFA for 24 hours. A small piece (5×5×5 mm) from an adjacent region was collected and immediately frozen in liquid nitrogen for future processing. The remainder of the tissue (up to 10×10×10 mm) was processed as described previously to prepare a single-cell suspension for flow cytometry and cell sorting (*47*). Briefly, lung tissue was infiltrated with 2 mg/ml collagenase D (Roche) and 0.2 mg/ml DNase I (Roche) in HBSS with Ca^2+^ and Mg^2+^ using syringe with 30G needle followed by incubation for 30µminutes at 37°C with mild agitation, followed by chopping with scissors and subsequent incubation for 15 minutes at 37°C in the same digestion buffer. The resulting single-cell suspension was filtered through a 40µm nylon cell strainer and erythrocytes were lysed using lysing buffer (Pharm Lyse, BD Biosciences). Automated cell counting was performed (Nexcelom K2 Cellometer with AO/PI reagent). Following live/dead staining with viability dye (Fixable Viability Dye eFluor 506, eBioscience), cells were incubated with a human Fc-blocking reagent (Miltenyi) for 5 minutes at 4°C and a mixture of fluorochrome-labeled antibodies for 30 minutes at 4°C. Cells were sorted at the Northwestern University Robert H. Lurie Comprehensive Cancer Center Flow Cytometry Core facility using a 100 μm nozzle and 40 psi pressure (Special Order Research Product FACSAria III instrument, BD Biosciences). Alveolar macrophages were identified using sequential gating as singlets/CD45^+^/live/CD15^-^/CD206^++^/CD169^+^/HLA-DR^+^/high autofluorescence; alveolar epithelial type II cells were identified as singlets/CD45–/CD31–/EpCam^+^/HLA-DR^+^. Between 50,000–300,000 cells were sorted into MACS buffer, immediately pelleted, and lysed in 350µl of lysis buffer (RLT Plus, Qiagen) supplemented with beta-mercaptoethanol. Lysates were stored at −80°C until RNA extraction.

### RNA isolation

Whole lung tissue was homogenized prior to RNA extraction (BeadBlaster 24, Benchmark Scientific) in the presence of lysis buffer (RLT Plus, Qiagen). Total RNA from whole lung tissue and sorted alveolar macrophages and alveolar type II cells was extracted (AllPrep DNA/RNA Mini Kit, Qiagen) and quality and quantity were assessed (TapeStation 4200, Agilent).

### Single-cell RNA-seq

A single-cell suspension was prepared as described above. Viability exceeded 80% as determined by AO/PI reagent staining. Single-cell 3’ RNA-seq libraries were prepared using Chromium Single Cell V2 Reagent Kit and Controller (10x Genomics). Libraries were assessed for quality (TapeStation 4200, Agilent) and then sequenced on NextSeq 500 or HiSeq 4000 instruments (Illumina), for mouse and human single-cell RNA-seq libraries, correspondingly.

Initial data processing was performed using the Cell Ranger version 2.0 pipeline (10x Genomics). Loupe Browser files for mouse and human datasets were generated using *aggregate* function in Cell Ranger pipeline with normalization on mapped reads and can be viewed using Single Cell Browser (10x Genomics). Post-processing, including filtering by number of genes expressed per cell, was performed using the Seurat package V2.2 and R 3.4 (*16, 48*). Briefly, gene counts from donor cells and counts from cells of patients with pulmonary fibrosis were aggregated into two separate Seurat objects, which were then aligned to each other using the canonical correlation analysis procedure (*15*). Following clustering and visualization with t-Distributed Stochastic Neighbor Embedding (t-SNE), initial clusters were subjected to inspection and merging based on similarity of marker genes and a function for measuring phylogenetic identity using BuildClusterTree in Seurat. Identification of cell clusters was performed on the final aligned object guided by marker genes. Differential gene expression analysis was performed for each cluster between cells from donors and from patients with pulmonary fibrosis. Heatmaps, t-SNE plots, violin plots, and dot plots were generated using Seurat. Functional enrichment with Gene Ontology (GO) Biological Processes was performed using GOrilla (database updated from Gene Ontology Consortium on December 12, 2017) with the top 1,000 upregulated genes with pulmonary fibrosis ordered by FDR q-value, unless there were fewer than 1,000 significantly upregulated genes (q-value < 0.05), in which case all significant genes were used, compared against all remaining detected genes as a background (*49, 50*). Gene Set Enrichment Analysis (GSEA) was performed using the GSEA 3.0 GSEA Preranked tool for enrichment of the Pulmonary Fibrosis gene set (obtained from Comparative Toxicogenomics Database using MESH term D011658 in November, 2017 via Harmonizome portal), with genes ordered by log difference in average expression (*17–19, 51*).

### Bulk RNA-seq

Libraries for RNA-seq from whole lung tissue were prepared from 100 ng of total RNA using a 3’ mRNA-sequencing library prep kit (QuantSeq FWD, Lexogen) on an automated liquid handling platform (Bravo, Agilent). Libraries from flow-sorted alveolar macrophages and alveolar type II cells were prepared from 50–100 ng of total RNA using a poly(A) enrichment module (NEBNext Ultra RNA, New England BioLabs). The resulting libraries were assessed (TapeStation 4200, Agilent), multiplexed, and sequenced as single-end 75 base pair reads (NextSeq 500, Illumina). For whole lung tissue RNA-seq analysis, demultiplexing was performed using bcl2fastq (version 2.17.1.14), trimming was performed using BBDuk, alignment was performed using STAR (version 2.52 with the hg38 assembly) (*52*), and raw read counts were generated using HTSeq (*53*). For alveolar macrophage and alveolar type II cell analysis, demultiplexing was performed using bcl2fastq (version 2.17.1.14), trimming was performed using trimmomatic (version 0.33) (*54*), alignment was performed using TopHat2 (version 2.1.0 with the hg38 assembly) (*55*), and raw read counts were generated using HTSeq. For all analyses, normalized tables of counts were generated from raw counts using the DESeq2 R package and the rlog transformation function (*56*). Filtering for low counts was performed by excluding all genes from which the minimum level of expression across all samples was lower than the average of the third quartile level of expression for all genes.

Functional enrichment with GO Biological Processes was performed using GOrilla with the top 500 upregulated genes and top 500 downregulated genes with fibrosis ordered by FDR q-value compared against all remaining detected genes as a background (*49, 50*). Gene Set Enrichment Analysis (GSEA) was performed using the GSEA 3.0 GSEAPreranked tool testing for enrichment of the Pulmonary Fibrosis gene set (obtained from Comparative Toxicogenomics Database using MESH term D011658 in November, 2017 via Harmonizome portal) with genes ordered by negative log FDR q-value (*17, 18, 51*).

### Fluorescence *in situ* hybridization

Multiplex fluorescent *in situ* hybridization was performed using RNAscope® (Advanced Cell Diagnostics (ACD), Newark, CA). Mouse lungs were inflated to 15 cm of H_2_O and fixed with 10% neutral buffered formalin (Thermo 9990244) for 24 h.

Lungs were paraffin embedded, and 5 µm tissue sections were mounted on Superfrost Plus slides (Fisher). Slides were baked for 1 h at 60°C, deparaffinized in xylene and dehydrated in 100% ethanol. Sections were treated with hydrogen peroxide (ACD 322381) for 10 min at room temperature, then heated to mild boil (98-102°C) in 1x target retrieval reagent buffer (ACD 322001) for 15 min. Protease Plus (ACD 322381) was applied to sections for 30 min at 40°C in HybEZ^TM^ Oven (ACD). Hybridization with target probes, preamplifier, amplifier, fluorescent labels and wash buffer (ACD 320058) was done according to ACD instructions for Multiplex Fluorescent Reagent Kit v2 (ACD 323100) and 4-Plex Ancillary Kit v2 (ACD 323120) when needed. Parallel sections were incubated with ACD positive (ACD 321811) and negative (ACD 321831) control probes. Sections were covered with ProLong Gold Antifade mountant (Thermo P36930). Probes used were: human *WNT7B* (ACD 421561, NM_058238.2) and human *AXIN2* (ACD 400241-C2, NM_004655.3); mouse *Wnt7a* (ACD 401121, NM_009527.3), mouse *Axin2* (ACD 400331-C3, NM_015732.4) and mouse *Wnt7b* (ACD 401131-C4, NM_009528.3). Opal fluorophores (Opal 520 (FP1487001KT), Opal 620 (FP1495001KT), Opal 690 (FP1497001KT), Perkin Elmer, Shelton, CT) were used at 1:1500 dilution in Multiplex TSA buffer (ACD 322809). Images were captured on Nikon A1R confocal microscope with 40x objective (NU-Nikon Cell Imaging Facility). Images were uniformly processed and pseudocolored in Fiji.

### Statistics

Statistical tools for each analysis are explicitly described with the results or detailed in figure legends.

### Supplementary Figures

**Supplementary Figure S1.**
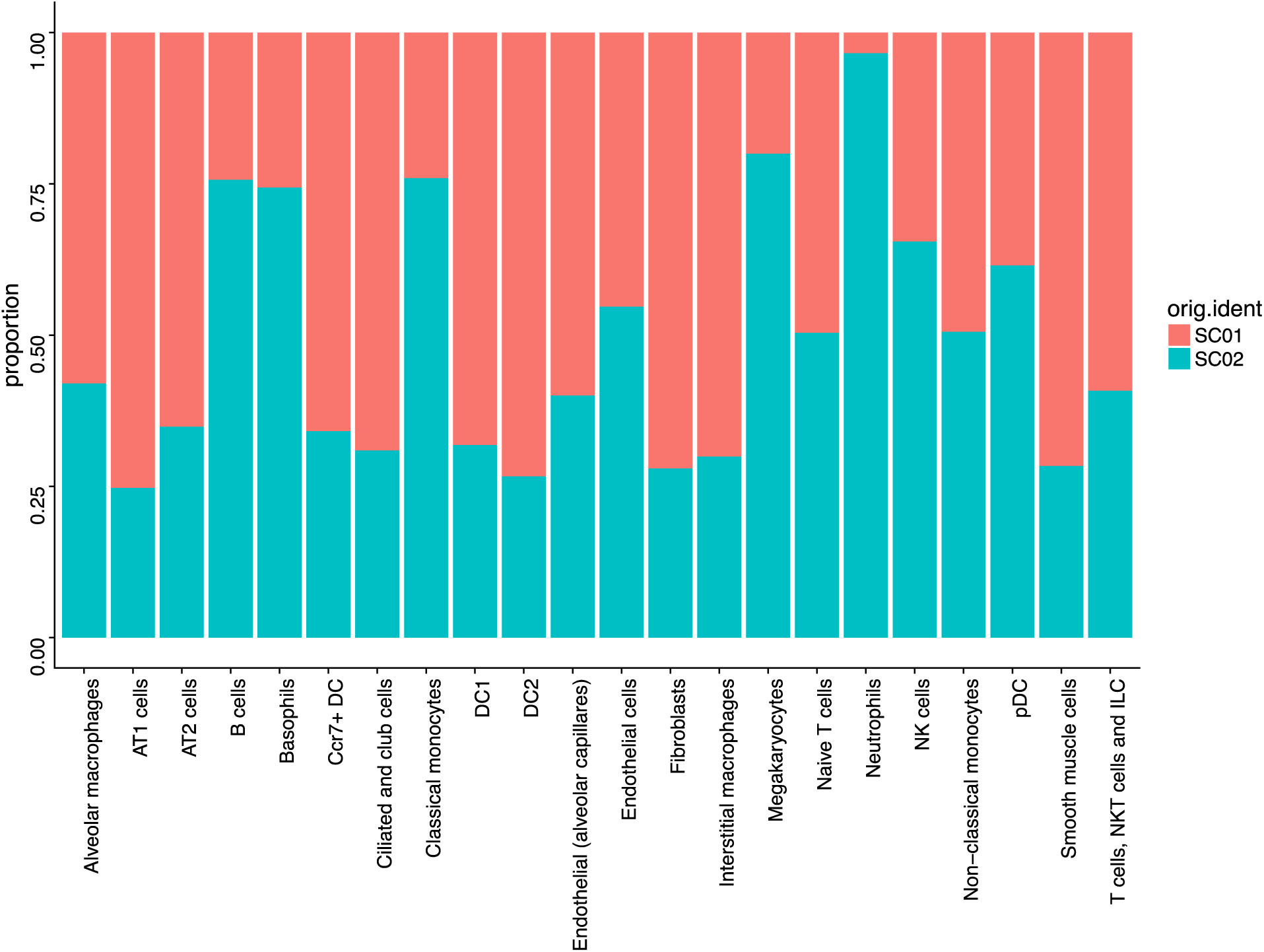
(Refers to Figure 1. Single-cell RNA-seq was performed on lungs from two mice and datasets were combined for analysis. Barplot demonstrates that all identified populations, including rare populations were well represented in both libraries.

**Supplementary Figure S2.**
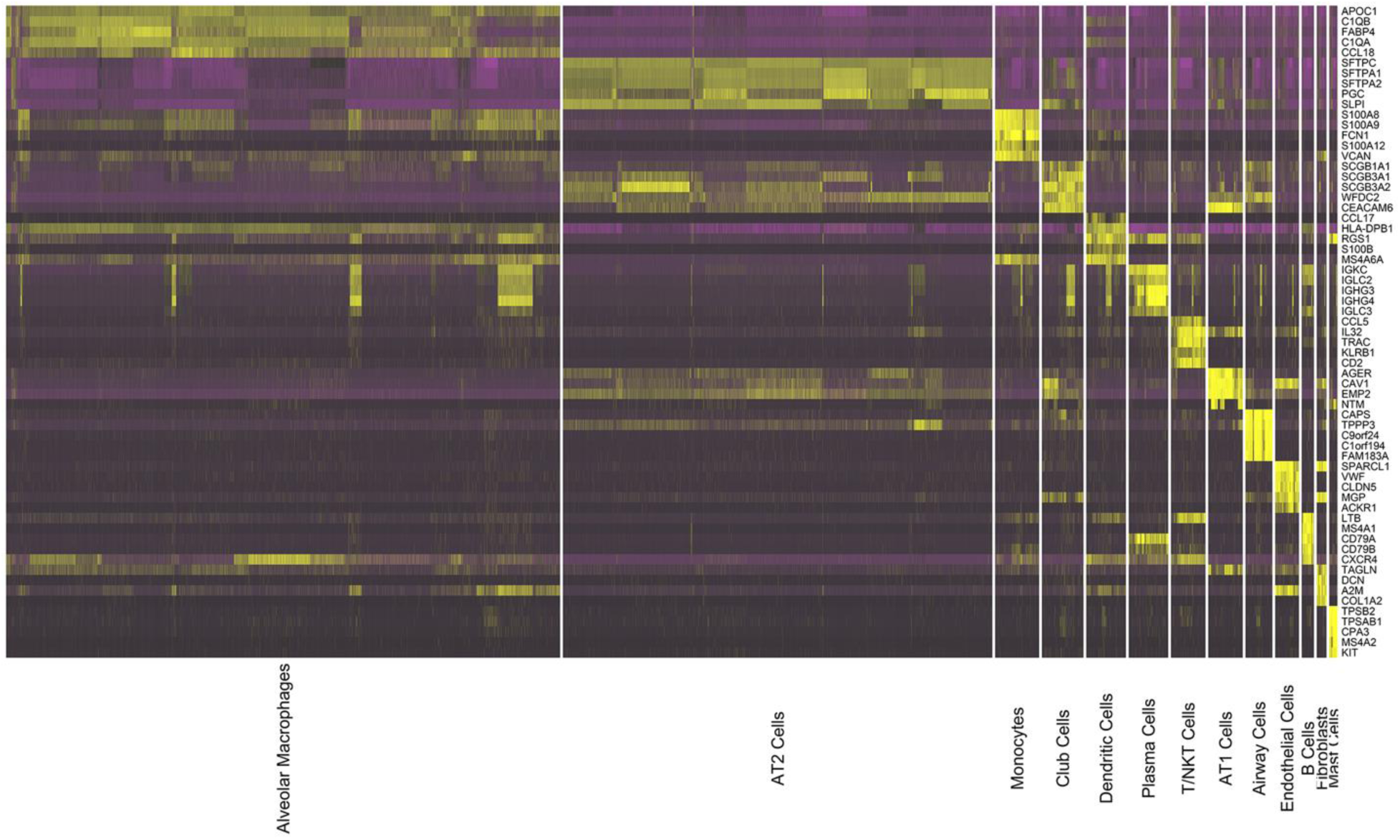
(Refers to Figure 2). Samples were obtained from six lung transplant donors, and six lung transplant recipients with pulmonary fibrosis. Single-cell suspensions were prepared and droplet-based single-cell RNA-seq was performed. Analysis of all twelve samples was performed using the R Seurat package with canonical correlation analysis procedure. Heatmap showing the top five positive marker genes within each cluster.

**Supplementary Figure S3.**
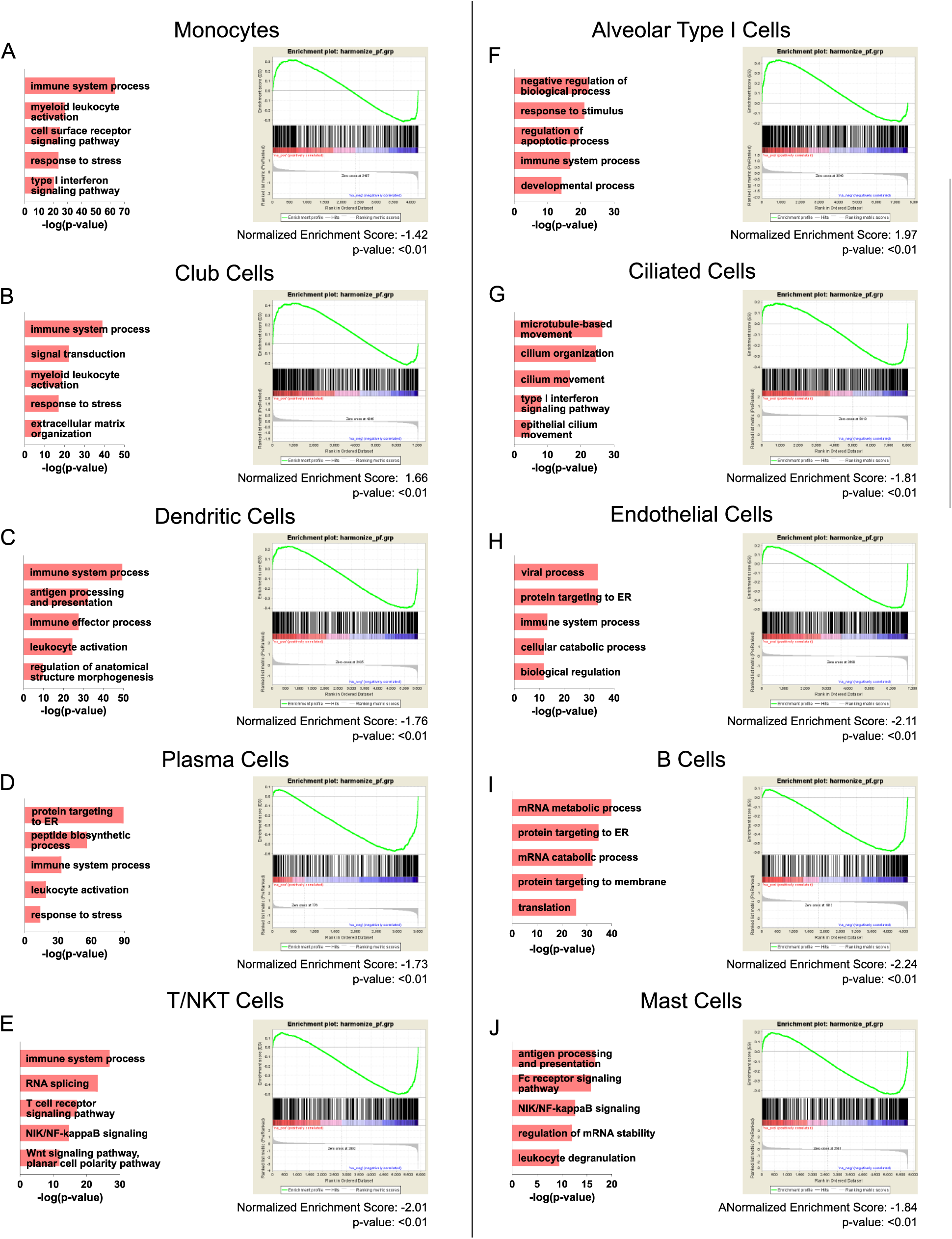
(refers to Figure 3). Representative significantly enriched GO processes and GSEA results using the Comparative Toxicogenomics Database Pulmonary Fibrosis Gene Set are shown for monocytes, club cells, dendritic cells, plasma cells, T/NKT cells, alveolar type I cells, ciliated cells, endothelial cells, B cells, and mast cells.

**Supplementary Figure S4.**
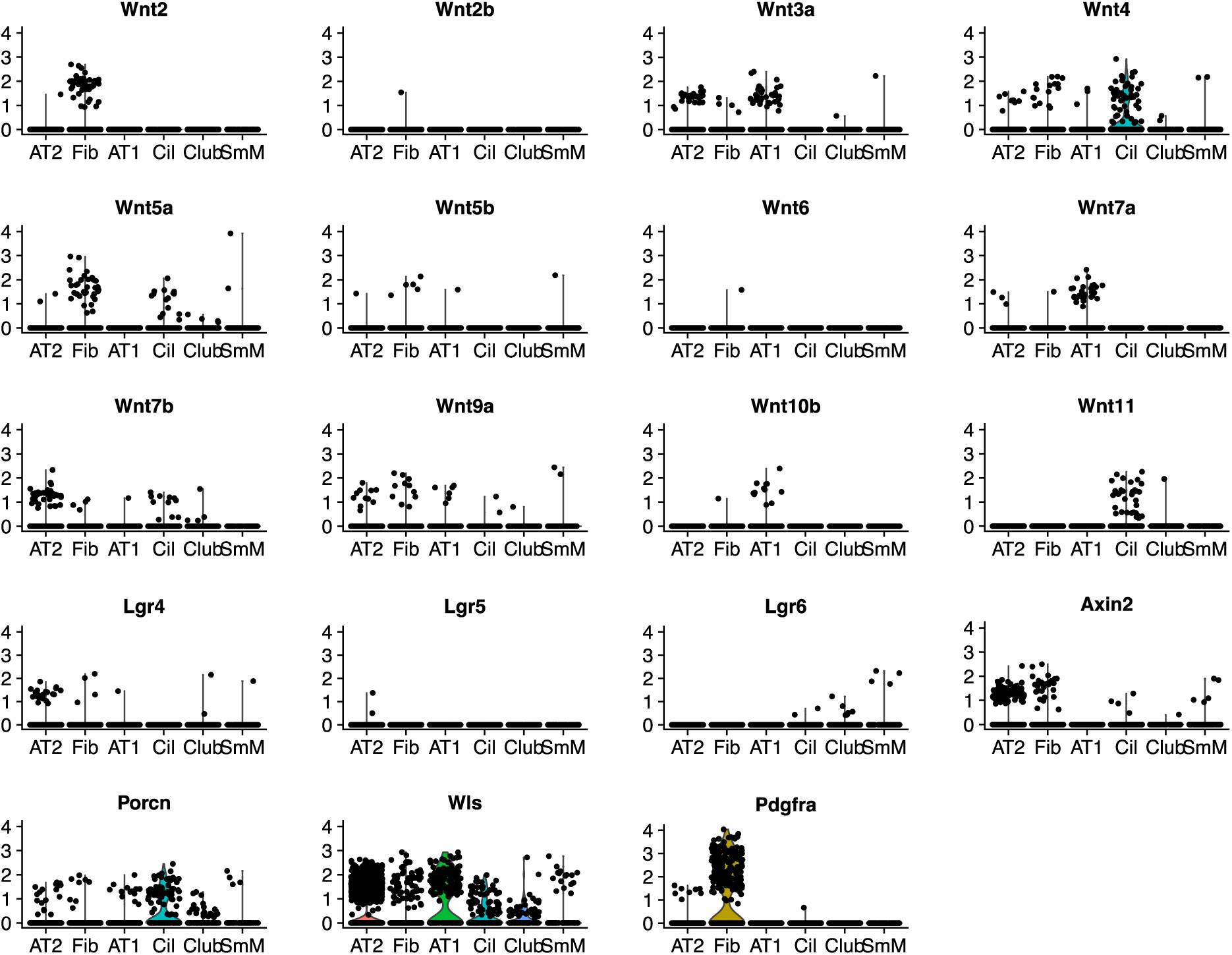
(refers to Figure 4). Violin plots demonstrating expression of the genes for Wnt ligands and related proteins reported by different groups to form niches responsible for maintenance of progenitor cell populations in mice in alveolar type I (AT1), alveolar type II (AT2), club cells (Club), ciliated cells (Cil), fibroblasts (Fib) and smooth muscle cells (SmM) (*21, 26, 41*). Expression of *Wnt1, Wnt3, Wnt8a, Wnt8b, Wnt9b, Wnt10a* and *Wnt16* were not detected and not shown.

**Supplementary Figure S5.**
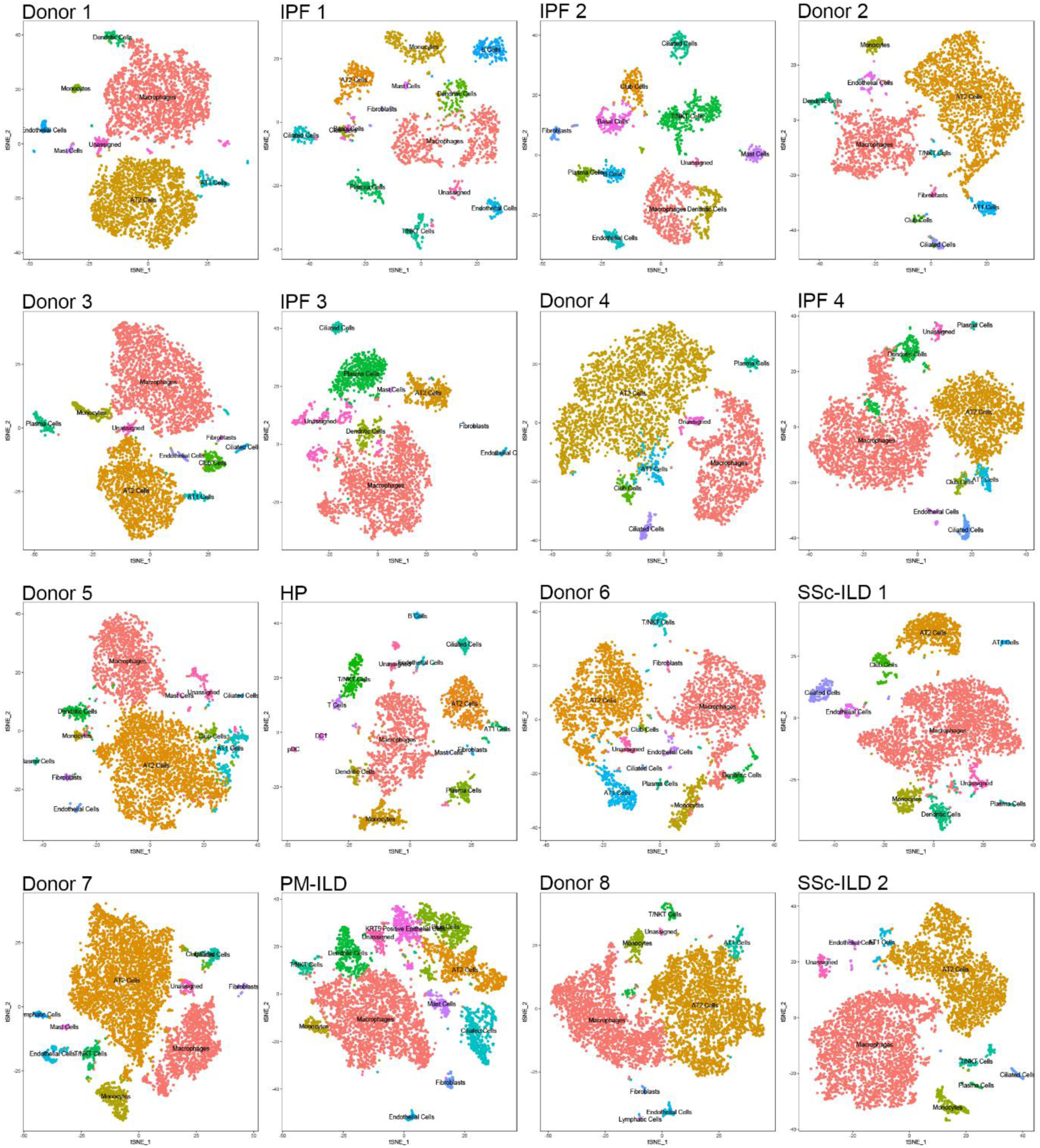
Individual t-SNE clustering of cells from 8 donor lungs and 8 patients with pulmonary fibrosis.

**Supplementary Figure S6.**
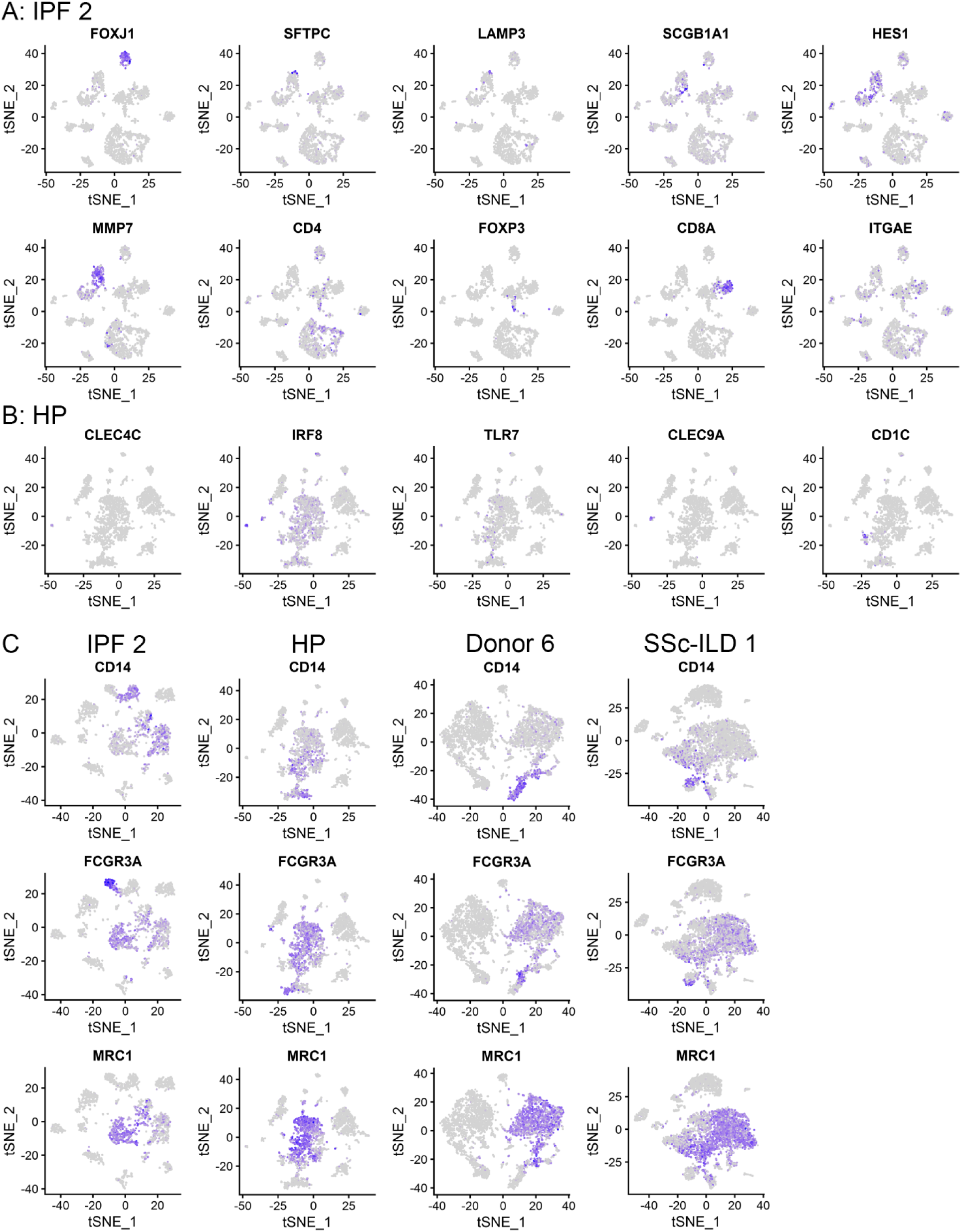
Examples of cell populations identified in a minority of patients. A. In subject IPF 2 there is a cluster of difficult to annotate. They were negative for the ciliated cell marker *FOXJ1* and highly expressed alveolar type II cell markers (*SFTPC*, *SFTPB*, *SFTPA2*, and *LAMP3*) as well as club cell markers (*SSCGB3A2*, *SCGB3A1*, and *SCGB1A1*). These cells also expressed high levels of *HES1*, *MMP7*. In this patient, *FOXP3* and *CD4* expressing regulatory T cells, and *ITGAE* and *CD8* expressing lung resident T cell subsets could be resolved. B. In the patient with hypersensitivity pneumonitis, two small populations of cells expressing CD45 were uniquely detected: One population expressed the plasmacytoid dendritic cell markers *CLEC4C*, *TCL1A*, *IRF8*, and *TLR7*, and the second expressed the conventional dendritic cell (DC1) marker *CLEC9A.* C. In four subjects (IPF 2, HP, Donor 6, SSc 1), classical and non-classical monocytes could be resolved based on expression of *CD14* and *CD16*. These cells do not express the macrophage marker *MRC1*.

**Supplementary Figure 7.**
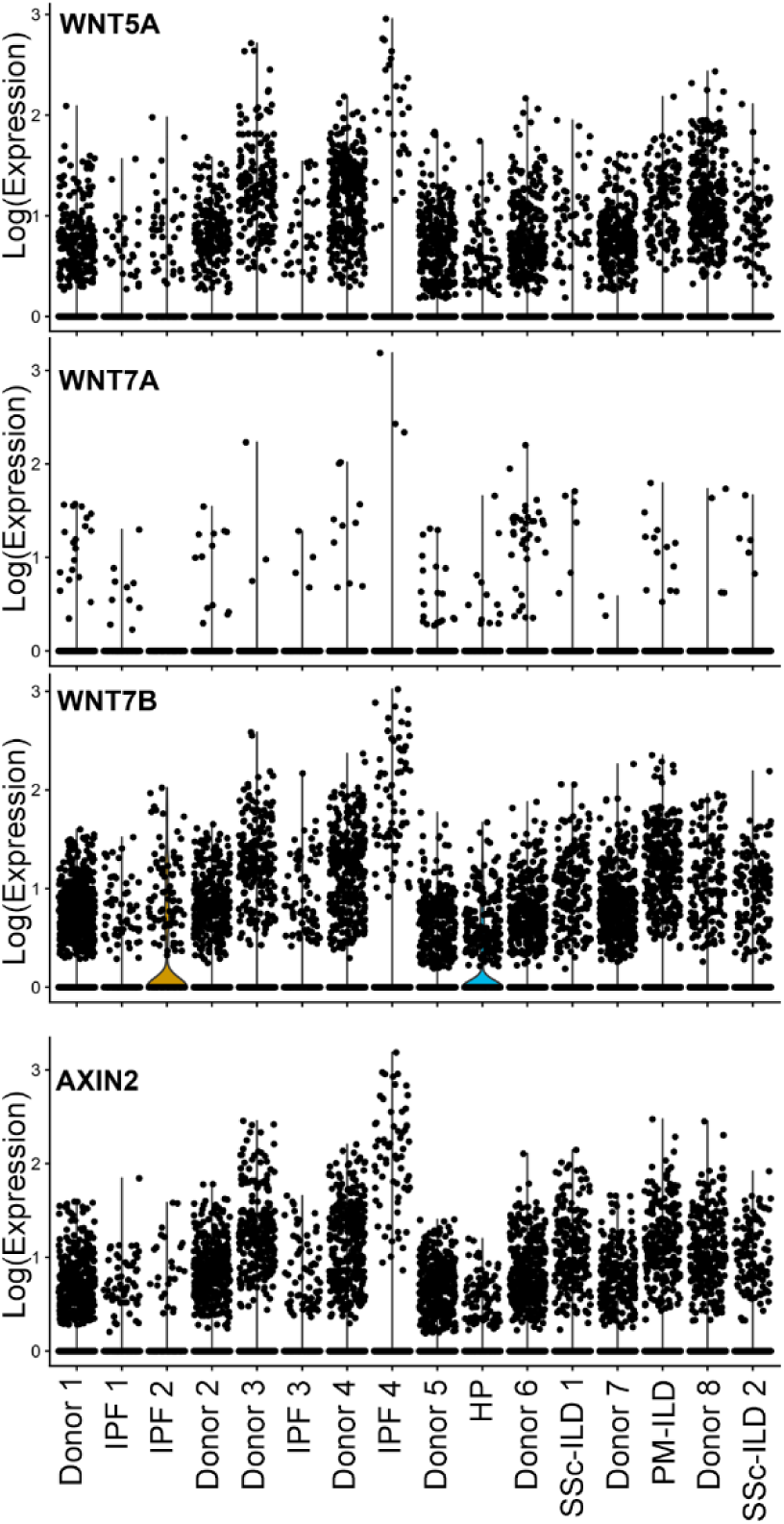
(refers to Figure 8). Wnt ligand expression is relatively homogenous in epithelial cells from the lungs of donors and patients with pulmonary fibrosis.

**Supplementary Figure S8.**
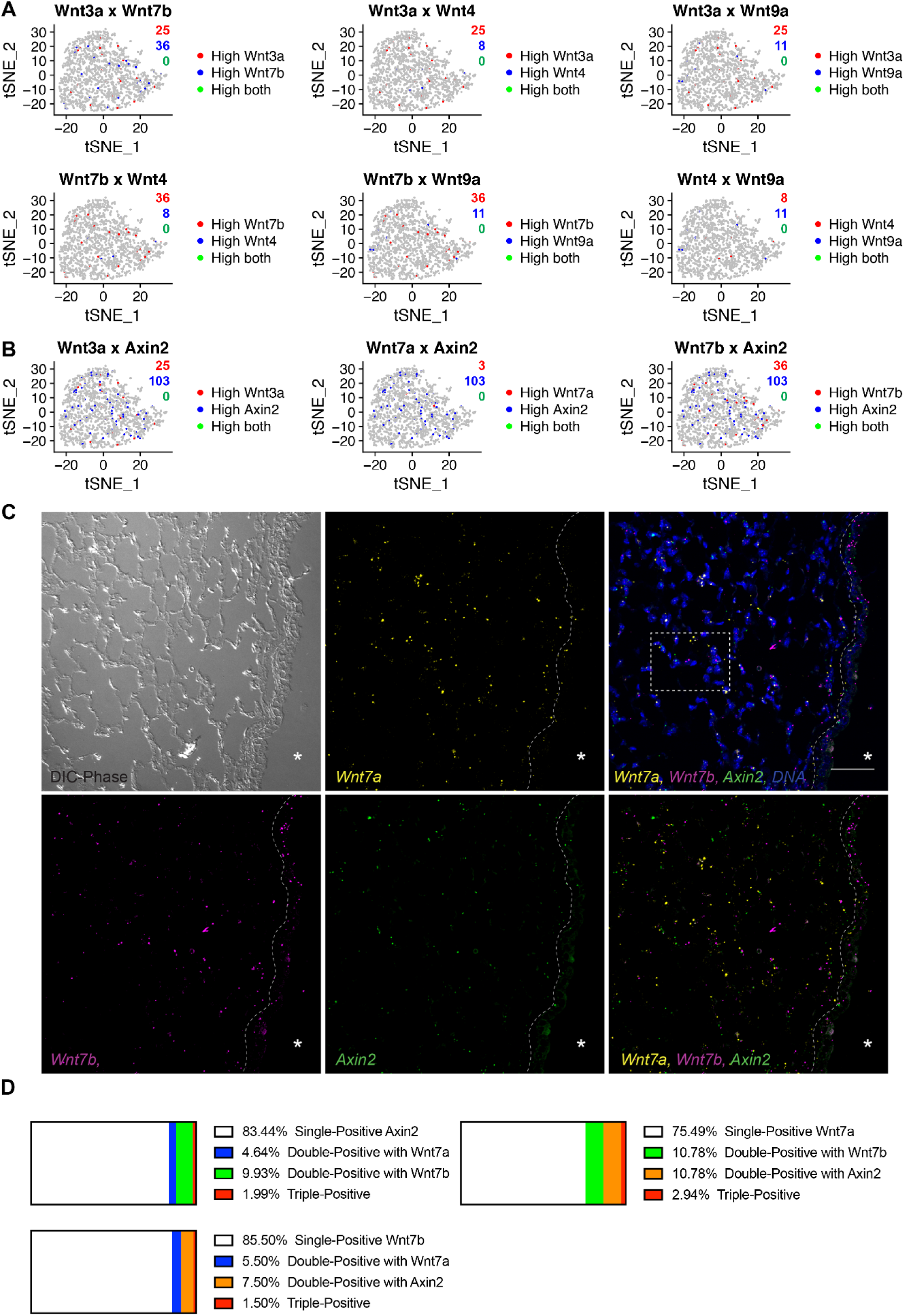
(refers to Figure 8). Mutually exclusive Wnt-expressor and - responder cells in the adult murine lung. A. Alveolar type II cells were examined for expression of individual Wnt ligands. *Wnt3a* and *Wnt7b* were the most abundantly expressed ligands, but no cells were detected that expressed more than one Wnt ligand. B. Cells expressing Wnt ligands were generally distinct from cells expressing the Wnt target gene *Axin2.* Clusters represent alveolar type II cells from two mice. C. Representative RNAscope in situ hybridization images of adult murine lung sections incubated with the designated target probes, *Wnt7b* (magenta), *Wnt7a* (yellow) and *Axin2* (green). Nuclei are stained with Hoechst (blue) and overlay images are shown on the far right. Differential interference contrast (DIC) phase image shown on left. As with human lung, *Wnt7b* is abundantly detected in airway epithelial/club cells (dotted line marks basement membrane of these cells). Boxed region corresponds to higher magnification view in Figure 8H. *Wnt7a* and *Axin2* are largely restricted to alveolar regions, where they mark distinct cells. Scale = 50µm. Asterisk (*) denotes airway lumen. D. Graph shows quantification of single-positive cells (*Wnt7b, Wnt7a* or *Axin2*; white), double-positive cells (with *Wnt7a,* orange; with *Wnt7b*, blue; with *Axin2*, green) and rare triple positive cells (gray) from 5 fields of view. Only signals directly overlapping with Hoechst-stained nuclei were counted, as cytoplasmic signals were difficult to assign to a particular cell. Double- and triple-positive cells were significantly less abundant than single-positive cells by ANOVA with Bonferroni’s Multiple Comparisons Test (p < 0.01). Clusters represent alveolar type II cells from two mice.

**Supplementary Figure S9.**
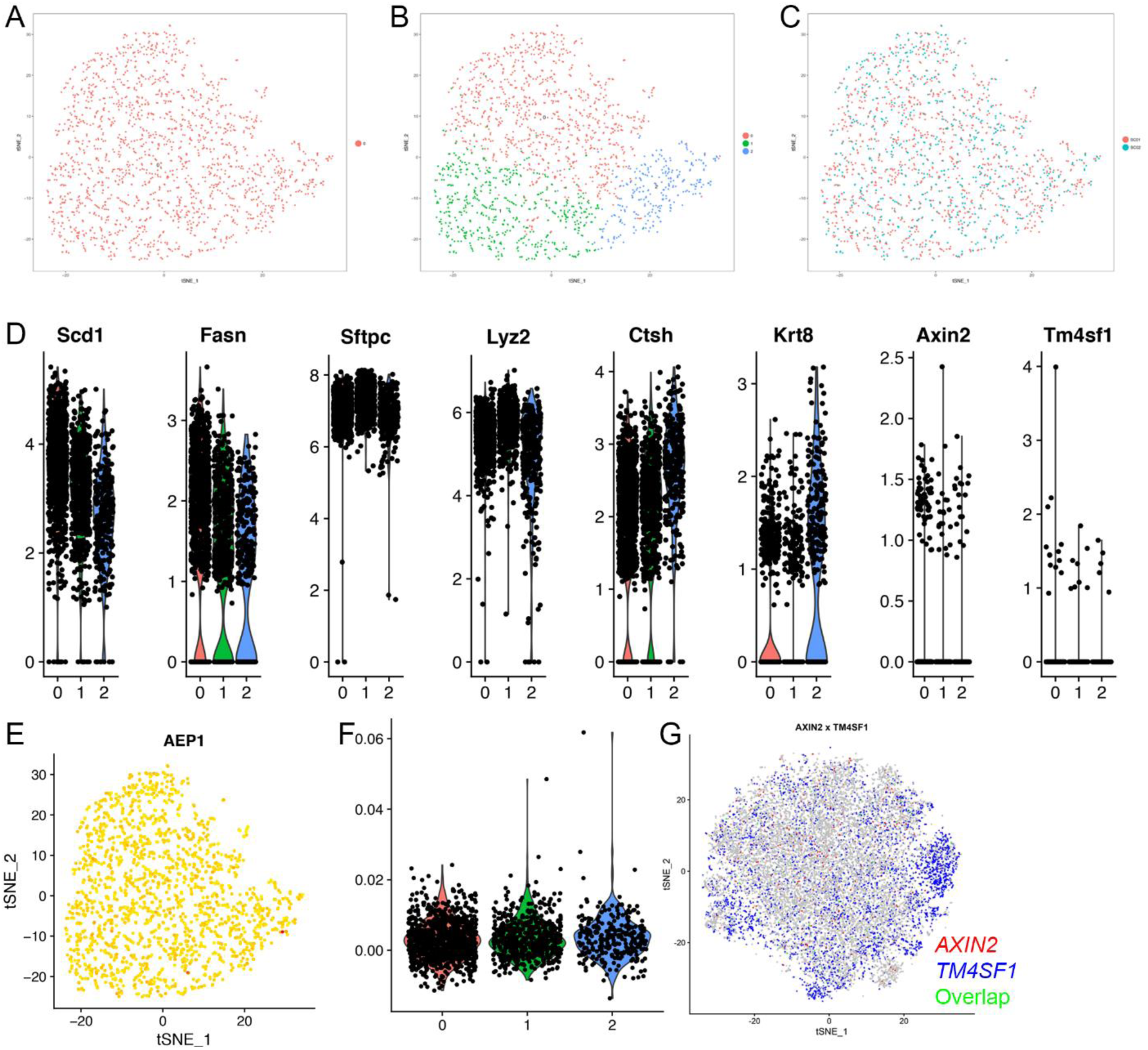
(refers to Figure 9).

**Supplementary Figure S10.**
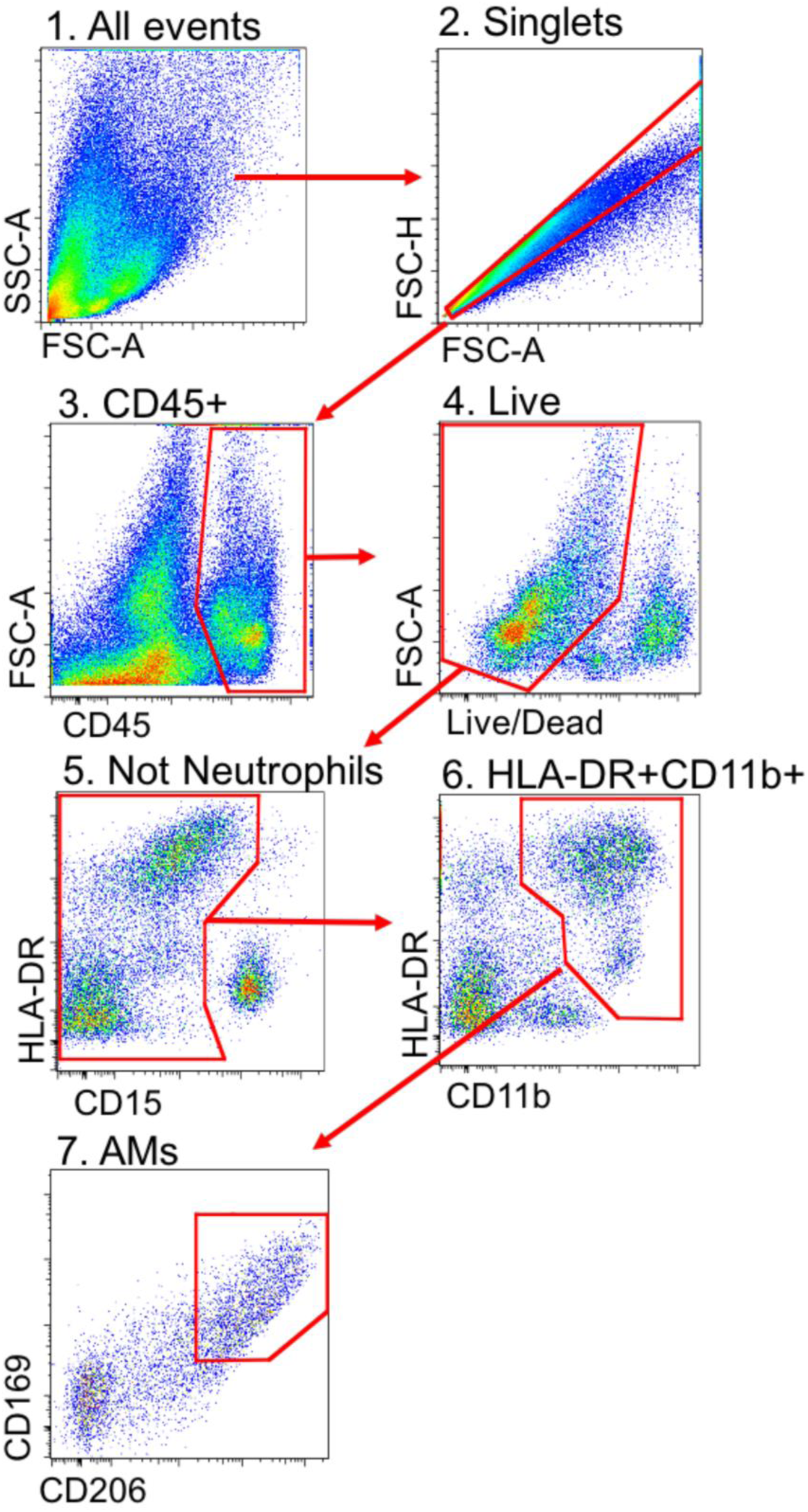
(refers to Figure 10). Gating strategy for flow cytometry performed on an *ex vivo* cryobiopsy.

### Supplementary Tables

Supplementary Table 1. Marker genes used to assign clusters from murine lung single cell RNA-seq. Refers to Figure 1.

Supplementary Table 2. Full list of identified differentially expressed marker genes in the CCA analysis. Refers to Figure 2.

Supplementary Table 3. Differentially expressed genes between donor and fibrosis lung samples in the 13 unique cell populations defined from the 12 patients (6 donors and 6 patients with lung fibrosis) in the CCA analysis. Refers to Figure 3.

Supplementary Table 4. Marker genes used to identify cell populations from individual t-SNE plots for each patient.

Supplementary Table 5. Full list of GO processes comparing gene expression in each macrophage cluster with average gene expression in alveolar macrophages.

## Acknowledgments

This work was supported by the Office of the Assistant Secretary of Defense for Health Affairs, through the Peer Reviewed Medical Research Program under Award W81XWH-15-1-0215 to Drs. Budinger and Misharin. Opinions, interpretations, conclusions and recommendations are those of the author and are not necessarily endorsed by the Department of Defense. Next generation sequencing on the Illumina HiSeq 4000 was performed by the NUSeq Core Facility, which is supported by the Northwestern University Center for Genetic Medicine, Feinberg School of Medicine, and Shared and Core Facilities of the University’s Office for Research.

Northwestern University Flow Cytometry Facility and Center For Advanced Microscopy are supported by NCI Cancer Center Support Grant P30 CA060553 awarded to the Robert H Lurie Comprehensive Cancer Center. Flow Cytometry Cell Sorting was performed on a BD FACSAria SORP system, purchased through the support of NIH 1S10OD011996-01. Multiphoton microscopy was performed on a Nikon A1R multiphoton microscope, acquired through the support of NIH 1S10OD010398-01. This research was supported in part through the computational resources and staff contributions provided by the Genomics Computing Cluster (Genomic Nodes on Quest) which is jointly supported by the Feinberg School of Medicine, the Center for Genetic Medicine, and Feinberg’s Department of Biochemistry and Molecular Genetics, the Office of the Provost, the Office for Research, and Northwestern Information Technology. The Genomics Computing Cluster is part of Quest, Northwestern University’s high-performance computing facility, with the purpose to advance research in genomics.

## Author contributions

PA Reyfman contributed to conceptualization, methodology, software, validation, formal analysis, investigation, data curation, writing, visualization, and supervision. JM Walter contributed to methodology, formal analysis, investigation, data curation, writing, and visualization. N Joshi contributed to methodology, formal analysis, investigation and writing. KR Anekalla contributed to methodology, software, formal analysis, data curation, and visualization. AC McQuattie-Pimentel contributed to methodology, validation, formal analysis, investigation, data curation, and writing. S Chiu contributed to methodology, investigation, formal analysis, and data curation. R Fernandez contributed to methodology, formal analysis, and data curation. M Akbarpour contributed to methodology, formal analysis, and data curation. C Chen contributed to methodology, validation, and investigation. Z Ren contributed to methodology, software, validation, formal analysis, and visualization. R Verma contributed to methodology, software, validation, and visualization. H Abdala-Valencia contributed to investigation, resources, data analysis, data curation, and project administration. K Nam contributed to methodology, investigation, and data curation. DR Winter contributed to methodology, software, validation, formal analysis, data curation, writing, and visualization. M Chi contributed to methodology, investigation, and data curation. S Han contributed to to methodology, investigation, and data curation. FJ Gonzalez-Gonzalez contributed to methodology and investigation. S. Soberanes contributed to methodology and investigation. S Watanabe contributed to methodology, validation, formal analysis and investigation. KJN Williams contributed to methodology, validation, formal analysis, investigation. AS Flozak contributed to methodology, validation, formal analysis, investigation and vizualization. TT Nicholson contributed to methodology and investigation. AC Argento contributed to methodology, validation, formal analysis, investigation, and data curation. CT Gillespie contributed to methodology, validation, formal analysis, investigation, and data curation. JD D’Amico contributed to methodology, validation, formal analysis, investigation, and data curation. M Jain contributed to conceptualization, data curation, writing, and supervision. BD Singer contributed to methodology, validation, formal analysis, writing, and visualization. KM Ridge contributed to conceptualization, resources, writing, supervision, and project administration. CJ Gottardi contributed to conceptualization, resources, writing, supervision, visualization and project administration. AP Lam contributed to methodology, validation, formal analysis, methodology, validation, formal analysis, investigation, and data curation. AV Yeldandi contributed to methodology, investigation, and formal analysis. VK Morgan contributed to methodology, validation, investigation, and data curation. M Hinchcliff contributed to resources, data curation, writing, and supervision. R Bag contributed to methodology, validation, formal analysis, investigation, and data curation. CL Hrusch contributed to methodology, validation, formal analysis, investigation, and data curation. RD Guzy contributed to methodology, validation, formal analysis, investigation, and data curation. CA Bonham contributed to methodology, validation, formal analysis, investigation, and data curation. AI Sperling contributed to conceptualization, methodology, validation, formal analysis, investigation, resources, data curation. RB Hamanaka contributed to methodology, validation, formal analysis, investigation, and data curation. GM Mutlu contributed to conceptualization, methodology, validation, formal analysis, investigation, resources, data curation, writing, visualization, supervision, and project administration. SA Marshall contributed to methodology and investigation. Luis Amaral contributed to conceptualization, software, writing, and supervision. A Shilatifard contributed to methodology and resources. H Perlman contributed to conceptualization, methodology, validation, formal analysis, investigation, resources, data curation, writing, visualization, supervision, project administration, and funding acquisition. JI Sznajder contributed to conceptualization, resources, data curation, writing, supervision, and project administration. A Bharat contributed to resources, data curation, writing, supervision, and project administration. SM Bhorade contributed to resources, data curation, writing, supervision, and project administration. GRS Budinger and AV Misharin contributed to conceptualization, methodology, validation, formal analysis, investigation, resources, data curation, writing, visualization, supervision, project administration, and funding acquisition.

## Funding

Paul A Reyfman is supported by Northwestern University’s Lung Sciences Training Program 5T32HL076139-13 and 1F32HL136111-01A1. James M Walter is supported by Northwestern University’s Lung Sciences Training Program 5T32HL076139-14 and Dixon Translational Research Grant. Deborah R. Winter is supported by the Northwestern Memorial Foundation Dixon Award, the Arthritis National Research Foundation, and the American Lung Association. SeungHyeHan is supported by Northwestern University’s Lung Sciences Training Program 5T32HL076139. Catherine Bonham is funded by K12 HL119995. Robert Hamanaka is supported by K01 AR066579 and ATS Foundation Unrestricted Grant. Robert Guzy is supporterd by K08 HL125910. Gokhan Mutlu is supported by NIH grants R01 ES015024, R21 ES025644 and U01 ES0236718. Ankit Bharat is supported by NIH grant HL125940 and matching funds from Thoracic Surgery Foundation, research grant from Society of University Surgeons and John H. Gibbon Jr. Research Scholarship from American Association of Thoracic Surgery. Benjamin D Singer is supported by HL128867 and the Parker B. Francis Research Opportunity Award. Jacob I Sznajder is supported by NIH grants AG049665, HL048129, HL071643, HL085534. Karen Ridge is supported by NIH grants HL079190 and HL124664.

Harris Perlman is supported by NIH grants AR050250, AR064546, AI092490, and HL108795, Funds provided by Solovy/Arthritis Research Society. GR Scott Budinger is supported by NIH grants ES013995, HL071643, AG049665, The Veterans Administration Grant BX000201 and Department of Defense grant PR141319. Cara J. Gottardi and Anna Lam are supported by NIH grant HL143800. Anna Lam is supported by NIH HL127245, Scleroderma Foundation and Respiratory Health Association grants. GR Scott Budinger is supported by NIH grants ES013995, HL071643, AG049665, The Veterans Administration Grant BX000201 and Department of Defense grant PR141319. Alexander V Misharin is supported by NIH grants HL135124 and AI135964, Department of Defense grant PR141319, and BD Bioscience Immunology Research Grant.

